# Specialization and adaptation in pollen sterol use by wild bees

**DOI:** 10.1101/2025.07.31.667884

**Authors:** Ellen C. Baker, Ellen Lamborn, Katie Berry, Geraldine A. Wright, Philip C. Stevenson

## Abstract

1. Sterols are essential metabolites in eukaryots, acting as stabilizing components of membranes and steroid hormone precursors. Unlike other animals, insects cannot biosynthesise sterols de novo so must acquire them from diet. Honeybees selectively uptake a subset of pollen sterols for use in their own tissues, rather than converting b-sitosterol or stigmasterol to cholesterol, as occurs in most insect herbivores because they do not have the chemical architecture to modify sterols like other insects. We do not know if this strategy is also employed by other bee species.
2. Here, we measured the sterol profiles of 56 species representing all families of UK bees to identify whether the same pollen sterols are used by solitary and eusocial bee species.
3. The Δ5 sterols 24-methylenecholesterol, isofucosterol and β-sitosterol which are common components of pollen sterolomes were also the main sterols found in all bee species. Campesterol and cholesterol were recorded almost universally in bees but in proportionately small amounts, despite their low abundance in pollen.
4. Our data suggest an ecological rather than phylogenetic driver of bee sterols because related species differed as a function of pollen diet. Generalist species contained more 24-methlyenecholesterol than their specialist congeners suggesting a trade-off between competition for food and desirable sterols. Asteraceae specialists have adapted to use pollen sterols that are not used by generalists, potentially gaining an advantage over these species. Despite the uptake of less common sterols by specialists, they still relied on similar sterols to other bee species. The dietary requirements of specialists could be fulfilled by pollen from non-host species, suggesting that while essential, sterols are not the primary driving force of pollen foraging choices.

## 1 Introduction

The co-evolution of bees and flowers has led to a highly inter-dependent relationship whereby many flowers are reliant on animal pollination and bees are fully dependent on flowers for their nutritional intake (Cappellari et al., 2016). The composition of pollen must therefore compromise between maintaining the fidelity of visiting pollinators but reducing losses of high-value pollen. As a result, the nutritional composition of pollen is highly diverse and variable in nutrient ratios, amino acid profiles, defense compounds and lipids including sterols (Wright et al., 2018; Zu et al., 2021; Furse et al., 2023a; Palmer-Young et al., 2019; Arnold et al., 2014).

Sterols are essential micronutrients serving key functions as membrane components and steroid hormone precursors in animals and plants. Diets that lack essential sterols can lead to poor brood production in honeybees (Moore et al., 2025). The pollen of many plant species contains sterols that are absent from animal tissues and scarce in terrestrial plant vegetative tissues such as 24-methylene cholesterol, isofucosterol, and desmosterol (Villette et al., 2014; Zu et al., 2021, Baker et al., *in review*). Honeybees have lost the capacity to both synthesise cholesterol and dealkylate longer chain phytosterols into cholesterol (Furse et al., 2023b). Instead, they have adapted to using longer chain phytosterols directly from their pollen food, in place of cholesterol (Herbert *et al*., 1980; Moore et al., 2025). Bogaert et al., (2025) has shown that 24-methylenecholesterol and isofucosterol and required by honey bees to produce offspring. This distinguishes bees from many other herbivorous insects, which maintain cholesterol rich membranes through modifying the sterols they consume from plants (Jing and Behmer, 2020).

A similar state of phytosterol-dominated tissues previously reported in honeybees is inferred but not evidenced to be present in other bee species (Feldlaufer *et al*., 1993; Jing and Behmer, 2020; Vanderplanck, Zerck, *et al*., 2020). And, given the diversity of pollen sterol profiles that have now been revealed, it is possible that other bee species may show distinct sterol profiles based on differences in diet representative of specific sterol requirements. Wild bee species have received considerably less research attention than the model eusocial species *Apis mellifera* and *Bombus terrestris*. As an ecologically and taxonomically diverse group, individual studies may not be applicable to bees in other families. Research on the sterol requirements of certain solitary bees has shown that they contain phytosterols in varying proportions, distinct from honeybees. Examples include an almost complete dominance of 24-methylenecholesterol in *Diadasia rinconis*, isofucosterol proportions three times higher in Meg*achile rotundata* than honey bees and β-sitosterol accounting for over 50% of total sterolic content in *Colletes cunicularius* (Svoboda and Lusby, 1986; Feldlaufer *et al*., 1993; Vanderplanck, Zerck, *et al*., 2020).

Wild bees provide an important pollination service for many wildflower species as well as contributing to pollination of some crops in the UK (Hutchinson *et al*., 2021). They also exhibit a range of dietary specialisations in pollen foraging (hereafter ‘lecty’); monolectic species are dependent on a single plant species for pollen intake, a strategy which is rare and only demonstrated by a small number of UK species (Falk and Lewington, 2015). Oligolectic species will collect pollen from multiple, closely related species which belong to a single genus or family. As a result of their specialist foraging, these species can be limited in distribution and emergence time. For instance *Melitta dimidiata* is confined to Salisbury plain in the south of England due to its need for *Onobrychis viciifolia* (Falk and Lewington, 2015). Astereaceae pollen is associated with oligolectic bee species in the UK and is rarely gathered by polylectic species such as honey bees. This is referred to as the ‘Asteraceae paradox’ (Müller and Kuhlmann, 2008; Praz, Müller and Dorn, 2008; Vanderplanck, Gilles, *et al*., 2020). Polylectic bee species collect pollen from multiple families and are considered generalist foragers. In the UK all bumble bee species, honey bees and some solitary bee species are polylectic (Falk and Lewington, 2015). The UK also contains a number of socially parasitic (kleptoparasitic) bee species, including six cuckoo bumble bee species which do not have any physiological adaptations for collecting pollen on their bodies and rely on host bee species to provision their young (Falk and Lewington, 2015).

There are over 270 bee species native to the UK, which range in their distribution and conservation status. The causes behind these differences in distribution have been attributed to climate and flower-richness, late emergence times and, in some cases, colony size (Goulson et al., 2005; National Biodiversity Data Centre, 2022). Bumble bees are often target taxa of seed mixes designed to increase floral resources in amenity and agricultural settings (Edwards and Jenner, 2018). However, species selection for these mixes is not based on an understanding of the nutritional value of the species to a range of different species (Williams and Lonsdorf, 2018). Here we sought to broaden our understanding of sterol requirements in bees by surveying a range of bee species representing a wide ecological and taxonomic breadth of species. We present the findings from two datasets, one dedicated to wild bee species and the other targeting bumblebees specifically. We examined the data for differences based on lecty and on phylogenetic relatedness and present a comprehensive overview of the trends in sterol requirements across these species.

## 2 MATERIALS AND METHODS

Two datasets were collected and analysed for this paper. The first was a set of bumblebee specimens, covering 18 species, compromising whole bodies, tagmata and pollen baskets. The second was a set of 56 wild bee species, comprising only whole-body samples. Processing, sample analysis and data analysis for both sets of samples is detailed below.

### 2.1 Sample collection

#### 2.1.1 Bumblebees

Female workers of all UK bumble bee species were targeted for collection. A minimum collection target of 5 individuals per species was set and specimens collected with landowner permission. Rare species (*Bombus distinguendus* and *B. sylvarum*) were collected only with prior arrangement with the Bumblebee Conservation Trust. Queens from abundant and common species (‘The big 7’: *Bombus hortorum*, *B. hypnorum*, *B. lapidarius*, *B. lucorum*, *B. pascuorum*, *B. pratorum* and *B. terrestris*) were collected to assess tagma composition. Female worker *Bombus lapidarius*, *B. terrestris* and *B. pascuorum* were collected from three different sites to facilitate intra-species analysis. For cuckoo species, which do not have workers, queens were collected, with males collected when this was not possible. For *Bombus ruderarius* and *B. campestris*, no female workers could be collected and so males was collected instead. To ensure there was not a difference between the sterols of male and females of the same species, males were opportunistically collected from well sampled common species (*Bombus hortorum*, *B. lapidarius*, *B. lucorum*, *B. ruderatus*, *B. rupestris*, *B. vestalis*) to enable statistical comparison.

Bees were captured using a net and euthanized in a -20°C freezer. They were then stored at -20°C and moved to -80°C storage in batches. After sample collection, bees were defrosted for dissection and the full gut, crop, sting and venom sacs removed from the abdominal cavity. This was to prevent misinterpretation of sterols detected from gut contents with those incorporated into body tissues. Any visible parasites were also removed. Ovaries, where present, were retained. A hind leg was also removed from all samples for future barcoding of specimens with indeterminate characters. For any male specimens, genitalia were retained, and guts removed. A set of queen bumble bees were also separated into body tagmata (head, thorax, abdomen). Any loose pollen in corbiculae or on the body was removed by brushing and full pollen baskets collected into 1.5ml glass vials. Dissected specimens were then refrozen at -80°C.

Species with indeterminate characters were barcoded to confirm identification as follows. Hind legs were stored at -80°C in a 1.5ml eppendorf. DNA was extracted from legs using a Qiagen DNeasy Blood & Tissue Kit following the supplementary protocol ‘Purification of total DNA from insects using the DNeasy® Blood & Tissue Kit’. Legs were homogenised using four 2.3mm ceramic beads in an MP FastPrep-24™ 5G bead beating grinder and lysis system in five cycles of 35s at 6 m/second followed by 180s rest. The genomic DNA was then sent to SureScreen Scientifics for PCR, barcoding and species identification.

#### 2.1.2 Wild bees

Species were chosen to cover a wide phylogenetic range and encompassed all foraging strategies (monolectic, oligolectic, polylectic and kleptoparasitic). Close relatives that had different foraging strategies were prioritised to understand the effects of phylogeny and ecology. A replication of three for each species was targeted initially but this was not possible for some species. Very common species were collected over three times, and from different sites to best capture intra-species variation. In total, 56 bee species were collected, with ten of these represented by a single sample. All samples were female except for *Melecta albifrons* which was represented by two male specimens. Collection was primarily focused in the South of England, from Royal Botanic Gardens (RBG), Kew and Wakehurst by HK. Other species procured from specific sites were done so with the permission of landowners. Bees were collected with a net then euthanised and stored at -20°C. Pollen was brushed from the outside of the body and removed from any pollen-collecting structures such as corbiculae or scopa. Guts were retained within specimens as removal would have likely damaged other internal tissues for the smallest specimens and so processing was kept consistent.

### 2.2 Sample preparation

#### 2.2.1 Bumblebees

All whole-body and tagma samples were covered in 5ml or 3ml of deionised water respectively and re-frozen at -80°C before being freeze dried. Samples were then transferred to plastic bead-beating vials containing four 2.3mm steel beads and homogenised at 30rpm in a Qiagen tissue ruptor for 3 minutes in two 1.5 minute intervals. Homogenised tissue was then transferred back to their glass vial in a 1:50 (mg:µl) dilution of GCTU and re-frozen at -80°C. For each sample, 60µl of homogenate was transferred to a 96-well plate for sterol extraction.

Paired pollen baskets weighing >20mg were separated and those weighing 10-20mg were combined for sterol analysis. Pollens were suspended in 200µl KOH and vortexed to disperse. They were then boiled at 75°C for two hours. Twelve which had dried out due to methanol evaporation during that time were re-suspended in 150 µl methanol. All saponified material was then transferred to a 96-well plate for sterol extraction.

#### 2.2.2 Wild bees

Sample processing was carried out by KB at RBG Kew. Specimens were covered in deionised water, frozen at -80°C and freeze dried until all water had been removed. Samples were then stored at -80°C before being homogenised. Specimens were homogenised in GCTU using a BioSpec Tissue Tearor with a 14mm head attachment. GCTU was made from Guanidine (6 M guanidinium chloride) and thiourea (1·5 M) dissolved in deionised H_2_O together and stored at room temperature out of direct sunlight.

### 2.3 Sample data collection

Samples were analysed in collaboration with staff at Royal Botanic Gardens (RBG), Kew. In addition to analysis samples, all plates contained samples designed to monitor instrument functioning: Blanks (60µl GCTU) were used to detect and remove background noise. All runs began with a blank. A stock solution of reference materials verified by multi-nuclear NMR (QA) were used to calculate coefficient of variance. Reference materials were available for: 24-methylenecholesterol, 24-methylenecycloartenol, anthelsterol (a sterol found in *C. anthelmintica*, ST(28:3), that is active under 330 nm UV but whose structure has not been determined formally using NMR), avenasterol, β-sitosterol, brassicasterol, campesterol, cholesterol, cycloartenol, cycloeucalenol, cyclolaudenol, desmosterol, episterol, ergosterol, isofucosterol, schottenol, sitostanol, spinasterol and stigmasterol.

The *m/z* and retention time for each sterol is shown in Table S5. Aliquots of 40 µl were used in the run. A QC solution of homogenised commercially available honey bee collected pollens (*Helianthus annus*, *Nymphaea* sp., *Fagopyrum esculentum*) and bumble bee adults, larvae and pupae in GCTU (6M guanidine + 1.5M thiourea to make 1 L dissolved in deionised water) was run at a series of concentrations to test the range of linearity of the instrument. Three concentrations of QC were used: 100%, 50%, 25%, corresponding to 40 µl, 20 µl and 10 µl aliquots of the QC stock solution. These were analysed in the same way as the samples to calculate the correlation between analyte concentration and signal size. Variables whose ratio was <0.75 were discarded.

All samples were then extracted as 96-well plates by SF (96-well plate, Esslab Plate+™, 2.4 ml/well, glass-coated) using a 96-channel pipette (liquid-handling robot, Integra Viaflow 96/384 channel pipette) as follows. First 150μl of internal standard (*d_7_*-cholesterol) was added to all samples. Concentration differed between plates as detailed above.

Then 500µl DMT (250µl ×2) was added to all samples. DMT was made from dichloromethane (DCM) (3 parts), methanol (1 part) and triethylammonium chloride (0.0005 parts, i.e. 500 mg/l), mixed and stored at room temperature out of direct sunlight. Followed by 500 µl deionised water before agitating using the multi-channel pipette. Layers were separated by spinning for 2 min (methanol + water, Solid, DMT + sterol) and 50 µl of extract (DMT & sterol) transferred to a 384-well plate and left for 30 min for the DMT to evaporate. Once the 384-well plate was full, 150 µl of LCMS quality methanol was added and the plate sealed with foil.

#### 2.3.1 Bumblebees

Sample run was prepared by EB, RTS and SF. Samples were run across two 384-well plates comprising eight individual 96-well plates. Sample order was randomised within each 96-well plate.

For plate A, whole-body bumble bee samples, the concentration of internal standard (*d_7_*-cholesterol) was 5 mg/l (1.99098e^−9^ Moles/sample). For plate C, body tagmata and pollen samples, it was 1 mg/l (3.98196e^−10^ Moles/sample).

#### 2.3.2 Wild bees

Sample run was prepared by EB, KB and SF. For each sample, 60µl of homogenate was transferred to a 96-well plate for sterol extraction. Samples were run across two 384-well plates comprising eight individual 96-well plates. Sample order was randomised within each 96-well plate. The concentration of *d_7_*-cholesterol for all plates in this run was 5 mg/l (1.99098e^−9^ Moles/sample).

### 2.4 Sample analysis and data processing

The sample run was completed under the supervision of SF following the protocol established in Furse *et al*. (2023a). Instrument output was also processed by SF as established in Furse *et al*. (2023a). Relative abundance was calculated by dividing the signal area for each metabolite by the signal of internal standard (*d_7_*-cholesterol).

These values were used for the majority of data analysis. For total sterol analysis, instrument output (*m/z*) was converted to mg/g first by dividing the signal area for each metabolite by the signal of internal standard (*d_7_*-cholesterol) then using the following formula:

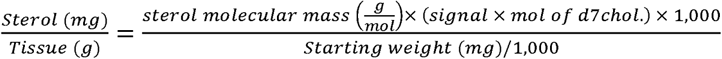

The amount of mol of *d_7_*-cholesterol was:

- Bumblebee plate (whole bodies): 1.99098e^−9^ Moles/sample
- Bumblebee plate (tagmata and pollen): 3.98196e^−10^ Moles/sample
- Wild bee plates (whole bodies): 1.99098e^−9^ Moles/sample

#### 2.4.1 Bumblebees

Due to the impact of the COVID-19 pandemic a second analysis run was needed to analyse samples of *Bombus distinguendus* and *B. jonellus* which were collected a year later than intended. Ten samples which had also shown anomalous results in the first run were also re-analysed as part of this run. The second analysis detected a different selection of low-abundance sterols compared to the previous run. As the primary aim of this work was to enable comparison between species and sites, any sterols which were not detected in both runs were therefore removed from data analysis to enable the most accurate comparisons. This led to the removal of two sterols from the second run (stigmasterol, ST(28:2)C, ST(31:2)A, ST(30:3)) and ten from the first (24-MCA, sitostanol, ST(27:2)C, ST(28:2)C, ST(29:0)A, ST(29:0)B, ST(29:2)A, ST(29:2)B, ST(29:2)C, ST(30:2)A, ST(30:2)C and ST(31:2)A). Proportions were re-calculated using only sterols detected in both runs (24-methylenecholesterol, β-sitosterol, campesterol, cholesterol, cycloartenol, cyclolaudenol, desmosterol, ergosterol and isofucosterol). Total sterol values for this dataset are therefore lower than if all detected sterols were included. Inclusion of this second data set did not alter the significance of any findings regarding phylogenetic and interspecies analysis.

#### 2.4.2 Phylogenetic analysis

Phylogenetic signal was calculated separately for each dataset following Zu *et al*. (2021). Phylo4d objects were created using phylo4d() from PHYLOBASE (R Hackathon et al., 2024).

Phylosignal(reps=999) from PHYLOSIGNAL (Keck *et al*., 2016) was used to calculate Pagel’s λ and Bloomberg’s K. These measures of phylogenetic signal were calculated for the individual proportions of all 19 sterols, total sterol (mg/g) and proportion of sterols belonging to different carbon chain lengths (27, 28, 29, 30, 31) and double bond saturation categories (cyclopropane ring(CPR), 0, 5, 7, NA). Ergosterol was classed as NA for double bond position as it has double bonds at both 5 and 7. All unidentified sterols were also classed as NA.

The phylogenetic trees were plotted by creating a phy-data object using treedata() from GEIGER v.2.0.11. (Pennell *et al*., 2014). Family and sub-genus clades were identified using getMRCA() from APE v.5.8.(Paradis and Schliep, 2019). Phylogeny was plotted with ggtree() from GGTREE v.3.12.0.(Yu *et al*., 2017, 2018; Yu, 2020). Heatmap and bar graph associated with phylogeny were plotted using ggplot() from TIDYVERSE v.2.0.0.(Wickham *et al*., 2019). The composite graph was then plotted using APLOT v.0.2.2.(Yu, 2023).

#### 2.4.3 Bumblebees

The most comprehensive phylogeny of the *Bombus* genus to date was used with permission from Hines (2008). Phylogeny was trimmed using keep.tip() from APE v.5.8.(Paradis and Schliep, 2019) to only include sampled species. *Bombus sylvestris* was not available in the phylogeny and so not included in these analyses. Subgenera were defined following Williams *et al*. (2008). Individuals of *Bombus terrestris* and the *B. lucorum* complex were combined into *Bombus terrestris*/*lucorum* for analysis as many samples had indeterminate characters. Similarly, *Bombus hortorum* and *B. ruderatus* were combined into *B. hortorum*/*ruderatus*. To plot these species groups on the phylogeny branch, tips for *Bombus terrestris* and *B. hortorum* were used.

#### 2.4.4 Wild bees

The supermatrix phylogeny produced by Henríquez-Piskulich, Hugall and Stuart-Fox (2024) was used to assess phylogenetic signal in the sterol dataset for the wild bee dataset. The 4,586 bee species dated chronogram with outgroups removed was selected for analysis. The tree was downloaded from Dryad, imported into R and pruned to 56 species using APE v.5.8.(Paradis and Schliep, 2019). Our dataset contained six out of the seven currently recognised families within the clade of Anthophila (bees): Andrenidae, Apidae, Colletidae, Halictidae, Megachilidae and Melittidae. Supplementary taxonomic data from Henríquez-Piskulich, Hugall and Stuart-Fox (2024) for all 56 species is available in the supplementary resources (Table S6).

### 2.5 Data analysis

All data analysis was done in R version 4.4.0 (2024-04-24 ucrt) (R Core Team, 2024) and R studio (2024.9.1.394) (Posit team, 2024). For each data type (whole-body, individual tagma, corbiculate pollen), a Bray-Curtis dissimilarity matrix was used to identify outliers using disana() from the LABDSV package v.2.1-0 (Roberts, 2023). Any data point with a minimum dissimilarity over 0.5 compared to all other points for that group was identified as an outlier, as per author guidance, and so was removed. No samples exceeded the outlier threshold in the bumblebee data, so all data was included in the analysis. For the wild bee dataset, each species with at least three replicates was tested for outliers. This led to the removal of five samples in total: One *Andrena flavipes*, two *Andrena florea*, one *Anthophora plumipes* and one *Colletes hederae*. Seven specimens were removed from total sterol analysis as their post-freeze drying weight was zero or negative; meaning their signals data could not be converted into mg/g units. This was likely due to incorrectly calibrated weighing equipment.

Summary statistics were calculated from species means using the STATS package from R (R Core Team, 2024). To carry out statistical comparisons of sterol profile between groups, proportion data (0-1) was used, where zero denotes a sterol is absent from a sample, and one denotes it is completely dominant. Data were arcsine square root transformed and used to calculate a Bray-Curtis dissimilarity matrix using vegdist(method=”bray”). The matrix was used to plot an NMDS using metaMDS(autotransform=FALSE). The lowest number of axes was selected where stress was ≤0.2 (established cut off, (Bakker, 2024)). The stress value reflects how well the ordination represents the original data. Adonis2() was used to carry out a PERMANOVA testing for significant differences between groups. Differences in variance between groups which could invalidate comparisons were tested using anova() and betadisper(). Pairwise comparisons between groups were carried out following Bakker (2024), using adonis2() with a Benjamini-Hochberg correction. All functions were from VEGAN (Oksanen *et al*., 2024) except anova() from the STATS package (R Core Team, 2023). A 2D biplot was created from NMDS with a stress value was <0.2. Strong (>0.7 absolute value) and significant (*p<*0.0100) sterol associations were plotted over the datapoints using ggplot() and ggrepel() from GGPLOT2 (Wickham, 2016) and GGREPEL (Slowikowski, 2024).

The INDICSPECIES package (De Cáceres and Legendre, 2009) was used to carry out Indicator Species Analysis (ISA) using multipatt() to determine significant associations between groups and individual sterols. ISA was done at the level of individual groups only and not for combinations of groups.

#### 2.5.1 Bumblebees

##### 2.5.1.1 Differences in sterol profile among UK bumblebee species

The six species with the highest number of samples, including male and female specimens, were used in an NMDS (trymax=200, k=3) to compare sterol profile between species. The first two dimensions of the NMDS are plotted in Figure S2. All available whole-body samples were used in an ISA to test for associations between individual sterols and species.

##### 2.5.1.2 Comparison between bumble bee collected pollen and whole-body tissue sterol profiles

A total of 40 pollen baskets large enough to analyse for their individual sterol content were collected from females. An NMDS was calculated using this data (n=40), all whole bee species (n=18) and all 295 pollen species detailed in Baker et al., *in prep* (n=295) (trymax = 200, k=2). ISA was then carried out to test for associations between these groups and individual sterols. As the pollen analysis had produced a wider range of sterols, only those present in both analyses were used to calculate total sterols and then proportions for the pollen data. These sterols were 24MC, β-Sitosterol, campesterol, cholesterol, cycloartenol, cyclolaudenol, desmosterol, ergosterol and isofucosterol.

##### 2.5.1.3 Intra-species sterol profile across different foraging environments

To determine whether there were significant intra-species differences driven by habitat, female workers of a set of species that are common in a range of habitats were collected from an urban garden environment (Royal Botanic Gardens, Kew), a semi-natural, dry, calcareous grassland site (sites near Rollestone Camp and Bulford Camp, Salisbury Plain Training Area, Wiltshire)(Natural England, 2015), and a semi-natural grass moor and heather moorland site (Harthope Valley, Cheviot hills, Northumberland National Park, Northumberland) (Natural England, 2013). These species were *Bombus terrestris/lucorum*, *B. pascuorum*, and *B. lapidarius*. Between site comparisons were plotted for *Bombus pascuorum*, *B. lapidarius* and *B. terrestris* individually using NMDS (trymax = 200, k = 2) and sites were compared statistically using PERMANOVA.

##### 2.5.1.4 Bumble bee body tagmata (head, thorax, abdomen) sterol profiles

Tagmata were analysed by NMDS (trymax = 200, k=2) and PERMANOVA to test for significant differences in sterol profile between head, thorax and abdomen. The strata argument was used to control for the effect of tagma being from the same individual. Species identity was also included as a fixed effect in the model. Additionally, ISA was carried out on the data to test for associations between tagmata and individual sterols.

##### 2.5.1.5 Sex differences in sterol profile

Where collection of females was not possible, male bumble bees were collected instead. For each species which had at least one male and female specimen (*B. barbutellus*, *B. hortorum/ruderatus*, *B. lapidarius*, *B. monticola*, *B. terrestris/lucorum*, *B. rupestris*, *B. sylvestris*, *B. vestalis*) an NMDS and PERMANOVA was calculated using means for each species-sex combination to prevent the often-higher female sample sizes affecting the results (trymax=200, k=2). Sex and species identity were included as fixed effects. *Bombus vestalis*, had highest number of male specimens (11) and a comparable number of female specimens (8) and was therefore used for a separate NMDS and PERMANOVA (trymax=200, k=2).

#### 2.5.2 Wild bees

##### 2.5.2.1 Differences in sterol profile across bee species

Species which were represented by at least 10 samples in the dataset were used for an NMDS comparison of inter-species differences. This included nine species from seven genera. An NMDS was calculated from proportion data using all available samples for these species (trymax=200, k=2).

##### 2.5.2.2 Differences in sterol profile between generalists and specialists

*Andrena* was the most speciose genus in the dataset and contained a similar number of oligolectic/monolectic (6) and polylectic (7) species which do not form monophyletic groups. *Andrena* species were therefore used in a NMDS (trymax=200, k=2) to calculate if the two lecty groups had significantly different sterol profiles.

##### 2.5.2.3 Differences in sterol profile between specialists

*Colletes hederae* and *C. succinctus* were both sampled from >3 sites with at least 10 replicates and have different foraging preferences. They were therefore used in an NMDS (trymax=200, k=2) to compare intra-species variability between two closely related species.

##### 2.5.2.4 Sterol profiles of monolectic and narrowly oligolectic bee species compared with their pollens

The dataset contained four monolectic bee species and two species which are narrowly specialised in the UK (*Colletes halophilus* and *Colletes hederae*). These bees were compared with their pollens and as bee-pollen groups with each other via NMDS (trymax=200, k=2) and PERMANOVA to determine whether specialist associations grouped separately.

##### 2.5.2.5 Asteraceae specialist bees

An NMDS (trymax=200, k=2) was calculated to compare the sterol profiles of solitary bees specialising on Asteraceae, non-Asteraceae specialists, polylectic bees and pollen from three tribes of Asteraceae (Carduoideae, Cichorioideae, Asteroideae). When comparing pollen and solitary bee sterol profiles, only sterols that had been detected in both datasets were used and proportions re-calculated using the remaining sterols. This led to a total of 17 sterols with ST(30:1)A and ST(30:1)B being removed from the bee dataset. Aligning data in this way will have altered the proportions of the remaining sterols in both datasets. However, to retain them would have made it impossible to fairly compare in an analysis.

Sterol data were then also grouped by B-ring double bond saturation (cyclopropane ring (CPR), Δ0, Δ5, Δ7, Δ8, NA) with ergosterol and all unidentified sterols classed as NA (ergosterol is a Δ5 and Δ7 sterol). In order to compare the proportions of different B-ring double bond groups between Asteraceae and non-Asteraceae specialist bees a Kruskal-Wallis rank sum test was carried out using kruskal.test() followed by pairwise Wilcoxon tests using pairwise.wilcox.tes(), both from the STATS package (R Core Team, 2024). Both analyses used a Benjamini-Hochberg correction for multiple testing.

## 3 RESULTS

### 3.1 The sterol profile of bees is largely defined by the presence of three pollen sterols

Total sterol content and sterol profile (sterolome) varied both within and between bee species (Figures 1 and 2). Nineteen sterols were detected across 56 wild bee species of which 13 were assigned using standards. The dataset included sterol analysis for seven eusocial *Bombus* species and 49 solitary bee species bee species and included 21 genera representing all bee families except for Strenotritidae which are restricted to Australia.

**Figure 1.**
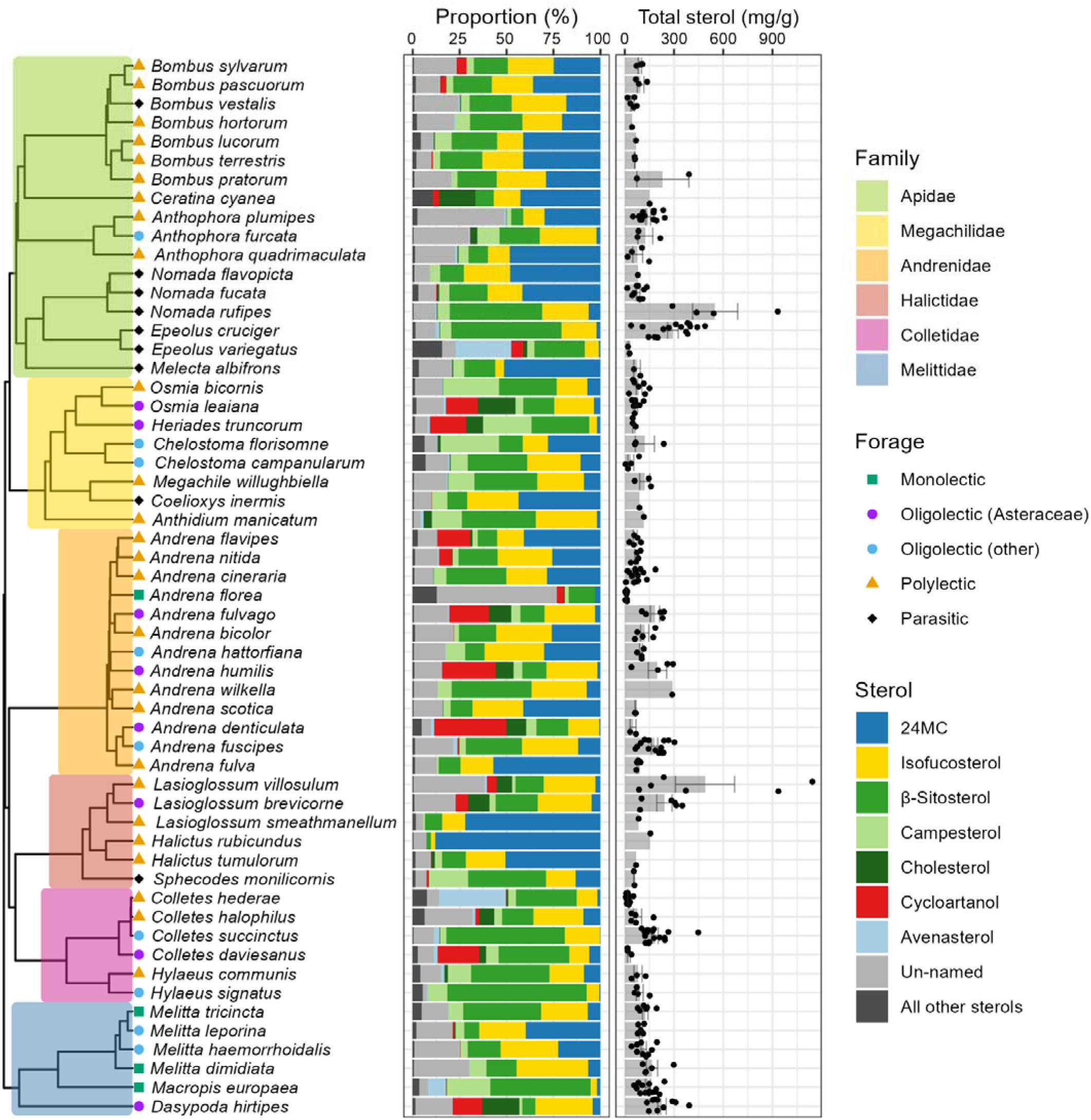
Phylogeny of 56 bee species subset from the supermatrix phylogeny produced by Henríquez-Piskulich, Hugall and Stuart-Fox (2024), showing family clades and foraging status. Species mean sterol percentages for seven key sterols are plotted along with total sterols (mg/g) for all samples (species means are shown by grey bars, error bars show ± standard error). 24-methylenecholesterol, isofucosterol and β-sitosterol account for the majority of sterols in most bee species. However, their proportions are highly variable between species. Bee families do not appear to show conserved sterolomes and only 3/19 sterols tested showed significant phylogenetic signal across the tree. Higher levels of cycloartanol appear across the tree and are often associated with species which are specialist (oligolectic) on Asteraceae flowers.

**Figure 2.**
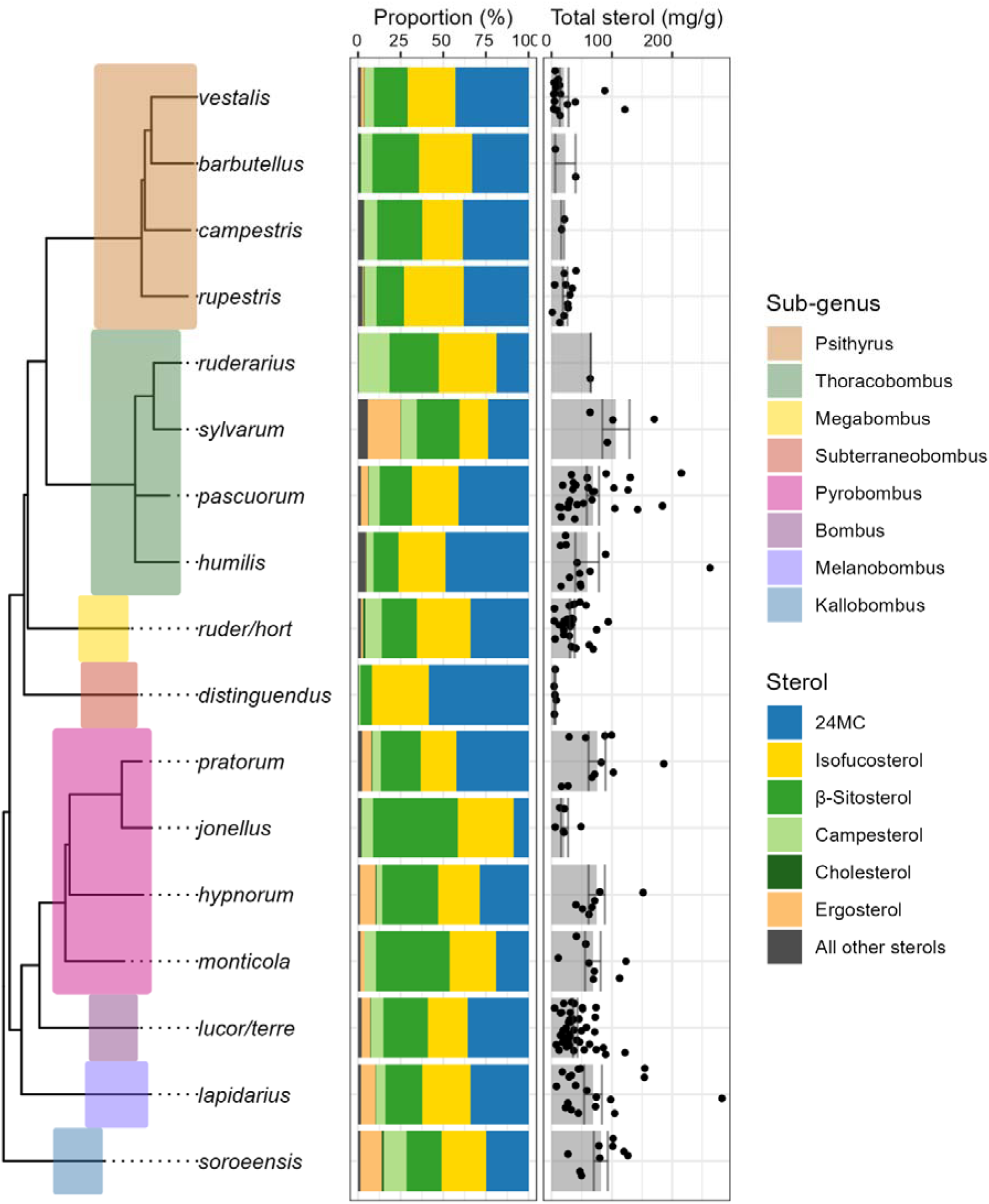
Species phylogeny subset from Hines (2008) with subgenera highlighted. Heatmap shows mean species proportion for the most commonly dominant five sterols plus cholesterol. Bar graph shows mean species total sterol production ±SEM. Points show individual samples of each species with different colours for each county the species was sampled from. Species which display a kleptoparasitic life strategy do not show a sterolome distinct from other Bombus species. All bumblebee species analused were dominated by 24-methylenecholesterol, isofucosterol and β-sitosterol, with no phylogenetic signal in any of the nine sterols analysed. Bombus sylvestris was omitted from this tree and so not included in phylogenetic analyses.

All bee sterolomes contained isofucosterol, β-sitosterol and 24-methylenecholesterol at a median percentage of >10%. The relative abundance of these three sterols varied as a function of bee species (Figure 1, Table S1). For example, the relative quantities of 24-methylenecholesterol within the genus, *Lasioglossum,* ranged from 3% in the shaggy furrow bee (*L. villosum*, species mean) to 72% in Smeathman’s Furrow Bee *L. smeathmanellum*.

The relative amount of β-Sitosterol was never lower than 2% and it was recorded < 10% for only six species. In addition, 24-methylenecholesterol and β-sitosterol were the only sterols that were recorded at >70% of the sterolome of any given species (e.g., β-sitosterol: *Hylaeus signatus*, 24-methylenecholesterol: *Halictus rubicundus*, *Lasioglossum smeathmanellum*). Fourteen of the nineteen sterols recorded accounted for less than 2% of the total sterols on average (Table S1). In Asteraceae specialists, cycloartanol was an important sterol (>15%: *Andrena denticulata*, *A. humilis*, *A. fulvago*, *Colletes daviesanus*, *Heriades truncorum*, *Osmia leaiana*, *Dasypoda hirtipes*) (Figure 1). The rarest sterols in the dataset were salisterol (26/56) and the fungal sterol, ergosterol (39/56) which may have been recorded as a result of fungal contamination of pollen (Table S1).

Most bee sterolomes contained at least 17 out of the 19 sterols we recorded (Table S3). Only three species contained <10 sterols, *Halictus rubicundus* (9), *Ceratina cyanea* (8), *Nomada flavopicta* (8). Five species contained no measurable cholesterol: *Andrena wilkella*, *Nomada flavopicta*, *Halictus rubicundus*, *Lasioglossum smeathmanellum* and *Coelioxys inermis*. Furthermore, campsterol was not detected in either *Halictus rubicundus* not *Ceratina cyanea*.

The proportions of three sterols in the bee sterolome showed phylogenetic signal (Figure 1 and 2): desmosterol (K=0.327, *p<*0.050, λ=0.790, *p<*0.005), campesterol (K=0.525, *p=*0.001, λ=1.012, *p=*0.001) and ST(28:1)B (K=0.479, *p<*0.050, λ=0.758, *p=*0.001) (Figure S1). Four additional sterols showed significance (i.e. *p* < 0.05) for at least one metric (Blomberg’s K or Pagel’s λ): ST(28:1)A, 24-methylenecholesterol, sitostanol, salisterol. We also grouped the sterols by carbon chain and B-ring saturation and found that the 28 carbon chain sterols group and Δ0 group showed significant phylogenetic signals for a single metric. In all except sitostanol, salisterol and the Δ0 group, this significance was in Blomberg’s K (Table S4).

We analysed separately the relationship between sterolome and phylogeny in the genus, *Bombus* (Figure 2). None of the sterols, carbon number/B-ring substitution or total sterol (mg/g) were associated with a significant phylogenetic signal in this genus. Across the bumble bee dataset (18 species), whole-body tissues were dominated by 24-methylenecholesterol, isofucosterol and β-sitosterol (Supplementary Table 2) and was similar to the sterolome of honey bees (Moore et al., 2025). Cholesterol was present in very low proportions across all species (< 0.5% median) compared to campesterol (median 6.6%). Ergosterol was detected in all species at a median proportion of 2.1%.

Total sterol quantities in the bees varied more than the sterol profile. For example, the shaggy furrow bee, *Lasioglossum villosulum,* showed the widest range with a maximum of 1143 ug/g and a minimum of 87 ug/g. The species with the highest mean values (*Nomada rufipes* = 551 ug/g and *Lasioglossum villosulum* = 489 mg/g) were both driven by outliers. For *Lasioglossum villosulum,* the two highest total sterol samples also had elevated quantities of the unassigned sterols, ST(28:1)B; these samples were collected from a different site than the other four samples. The red-thighed epeolus, *Epeolus cruciger,* had a less variable but higher total sterol content (293 ug/g) and was also better sampled.

### 3.2 The sterolomes of solitary bees differ significantly

In the nine most sampled bee species in our dataset, the sterolome was significantly different between every species except one of the kleptoparasites and its host (Figure 3A, PERMANOVA, F_8,118_=25.306, *p=*0.001, NMDS stress value = 0.151, pairwise comparisons, p < 0.05). In this case, the ashy mining bee, *Andrena cineraria,* and its kleptoparasite, *Nomada fucata*, had a distinct sterolome from other bee species but the two species were not significantly different from each other (*p>*0.100). It is notable that the sterolome of the heather colletes, *Colletes succinctus,* and its kleptoparasite, *Epeolus cruciger,* shared some sterols presumably due to their parasite-host relationship but nevertheless still differed significantly.

**Figure 3.**
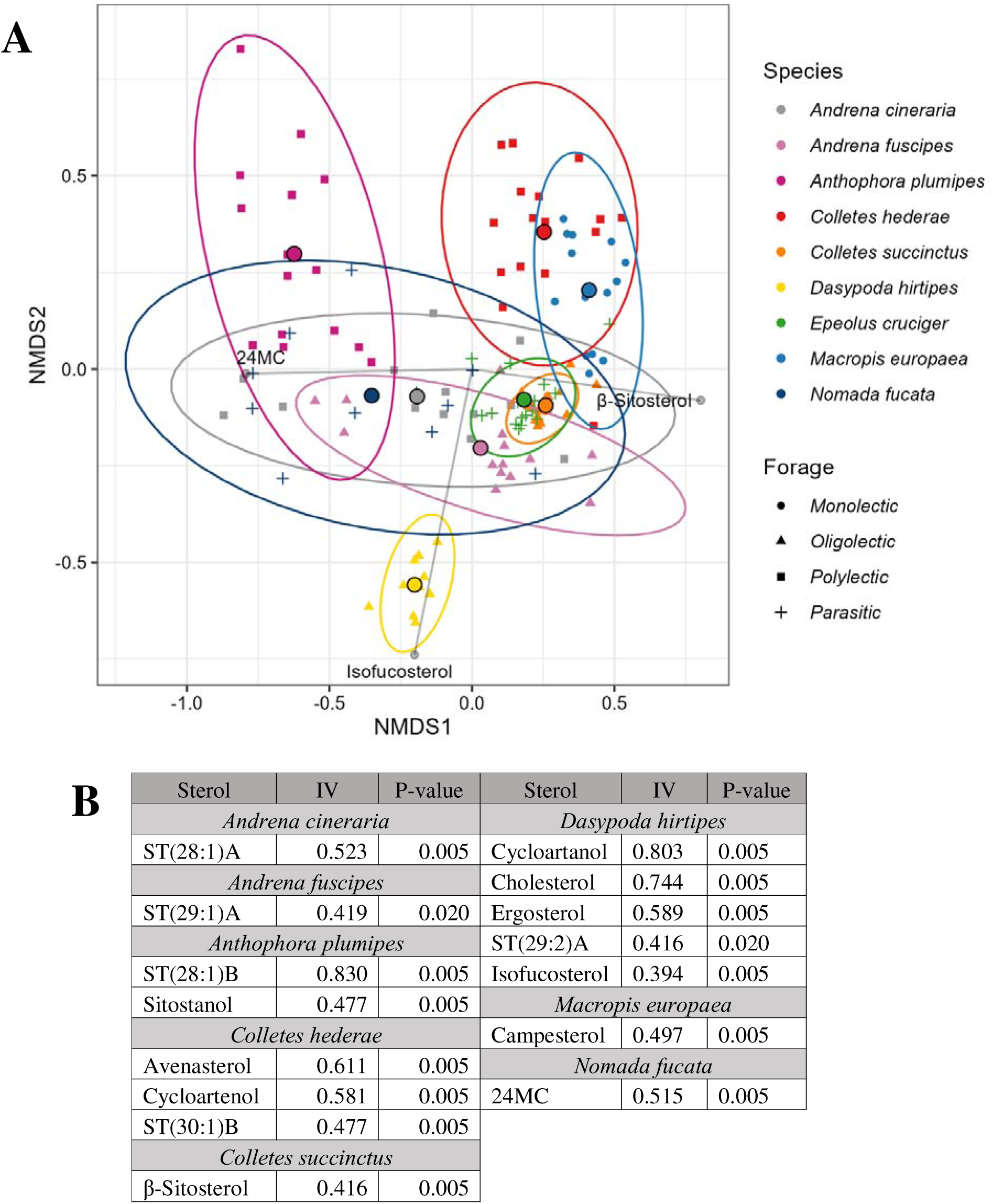
**A)** Interspecies comparison of sterol profile among species with ≥ 10 samples demonstrating significant interspecies differences in sterolome (PERMANOVA, F_8,118_=25.306, p=0.001). Grey lines indicate significant (p<0.010) and strong (absolute value >0.7) correlations. There are clear similarities in the sterol profile of ecologically linked species such as Epeolus cruciger with Colletes succinctus and Nomada fucata with Andrena cineraria, both of which are kleptoparasite-host pairs. All pairwise comparisons were significantly different (p < 0.05) except Nomada fucata and Andrena cineraria. There were significant differences in variance between species with Colletes succinctus displaying the most conserved sterolome and Nomada fucata the most variable. NMDS stress value = 0.151. **B)** Indicator Species Analysis of species with ≥ 10 samples as shown in Figure 3A. Of the 19 sterols tested, 15 were indicators of individual species groups. Indicator values (IV) and p-values are shown for significant sterols. Dasypoda hirtipes was associated with the most sterols and shows a distinct grouping in Figure 3A. Epeolus cruciger was not associated with any individual sterols, potentially as a result of its high similarity in sterolome with Colletes succinctus as well as other bee species.

Species identity explained over 50% of the variation in the data (R^2^=0.632). However, species groups also displayed significantly different variances (Fig 3, F_8,118_=6.825, *p<*0.001). All species except *Epeolus cruciger* were associated with a difference in at least one sterol (Figure 3B). This may be due to the strong similarity in sterolome between *Epeolus cruciger* and *Colletes hederae* which was associated with three sterols (Figure 3B). The pantaloon bee, *Dasypoda hirtipes,* an Asteraceae specialist, showed the most distinct sterolome which included strong associations with cycloartanol and cholesterol (Figure 3B).

In comparison to the larger dataset, the sterolomes of bumblebees exhibited only small differences in the proportions of sterols (Figure S2, PERMANOVA: F_5,145_=2.938, *p*=0.001, NMDS stress value = 0.128). The six most sampled bumblebee species showed no significant difference in the sterolome (F_5,145_=2.095, *p*>0.050). Species identity explained a low proportion of the variation in the data (R^2^=0.092). Pairwise comparisons showed the garden/ruderal bumblebee, *Bombus hortorum/ruderatus* and the brown-banded carder bee, *B. humilis*, had the most distinct sterolomes. Both differed significantly from *B. pascuorum* (*p*<0.050) and *B. terrestris/lucorum* (*p*<0.050) as well as each other (*p*<0.050). In addition, *B. humilis* differed significantly from the red-tailed bumblebee, *B. lapidarius* (*p*<0.050).

Comparing all bumblebee species, the red-shanked carder bee, *B. ruderarius,* was more associated with campesterol (IV=0.32, p<0.050). It is worth noting the indicator value for this association is very low (<0.4) suggesting there is little difference between species that can be attributed to individual sterols. Furthermore, *B. ruderarius* was represented only by a single sample.

### 3.3 Generalist and specialist bees show subtle differences in their sterolome

While all bumble bees (*Bombus*) are generalist foragers, other genera display a wider variety of foraging types. For instance, *Andrena* contains polylectic, oligolectic and monolectic species. Generalist *Andrena* species (polylectic) showed a significantly different sterol profile to specialist *Andrena* (oligo/monolectic) (Figure 4, NMDS: stress = 0.077, PERMANOVA: F_1,11_=3.417, *p<*0.050, dispersion: F_1,11_=1.105, *p>*0.100). Generalist bees in the genus *Andrena* spp. were strongly and significantly associated with 24-methylenecholesterol (IV= 0.85, *p=*0.010) and the unassigned sterol ST(28:1)B (IV=0.86, *p<*0.015). In contrast, specialist species were associated with cholesterol (IV=0.924, *p*<0.015) and the unidentified sterol ST(30:1)A (IV=0.76, *p*=0.025). However, foraging strategy only explained a small proportion of the variation in the data (R^2^=0.237).

**Figure 4.**
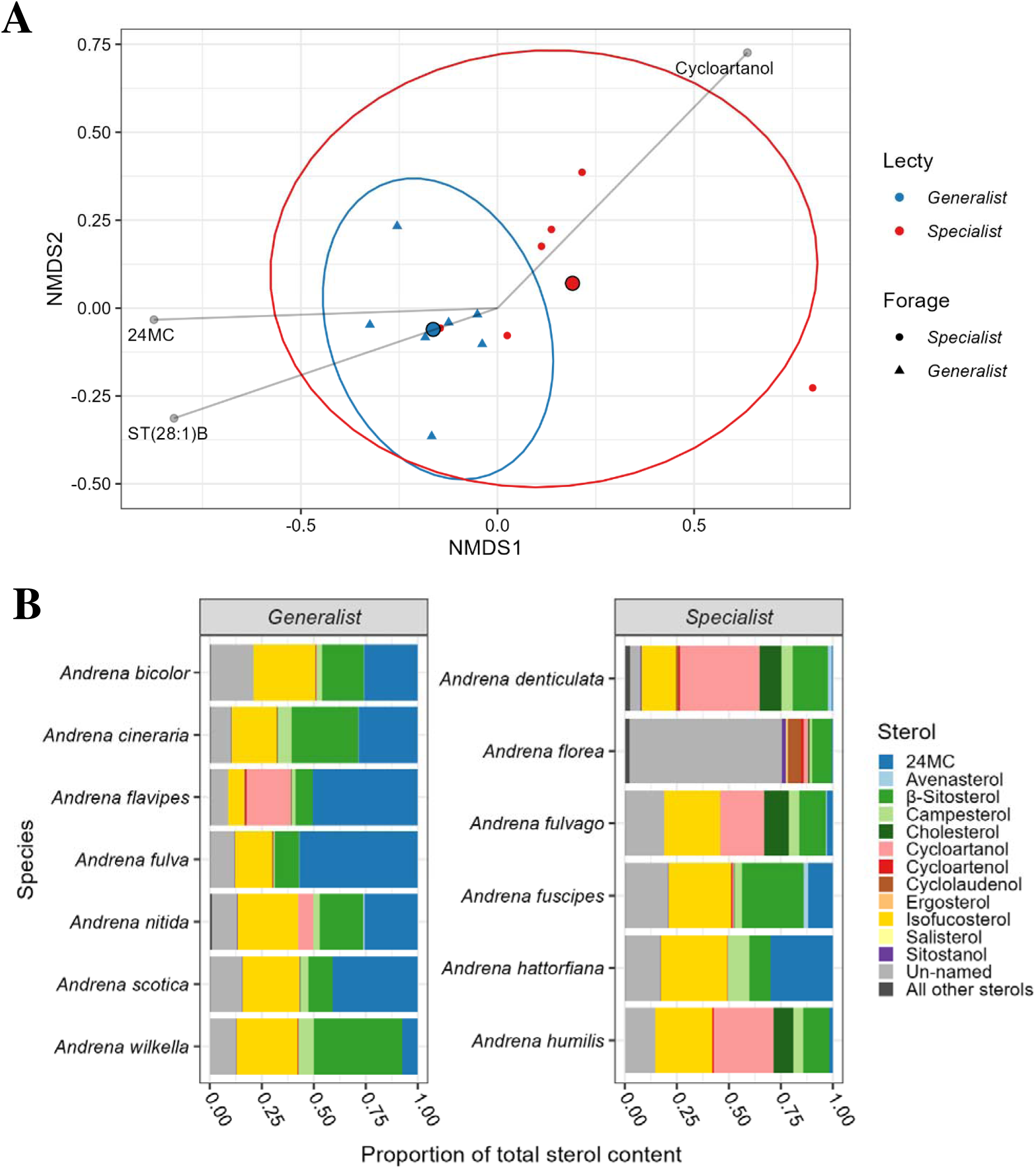
**A)** NMDS of generalist (polylectic) and specialist (oligolectic/monolectic) Andrena species (stress value =0.077) which showed a significant difference between groups (PERMANOVA: F_1,11_=3.417, p<0.050). Grey lines indicate significant (p<0.010) and strong correlations (absolute value >0.7). Generalist species were correlated with higher 24-methylenecholesterol and specialist species are associated with higher cholesterol. **B)** Proportions of dominant sterols across all Andrena species included in Figure 6A. All remaining sterols are collapsed into a single group. Generalist species show higher levels of 24-methylenecholesterol and specialist species that target Asteraceae species (A. denticulata, A. fulvago and A. humilis) show higher proportions of cycloartanol. Andrena florea contains very high proportions of un-named sterols suggesting this bees sterolome requires further characterisation.

In addition to differences between generalist and specialist bees, it should also be expected that specialist bees displayed distinct sterol profiles as a result of their host plant pollen. Two specialist bees in the genus, *Colletes,* illustrate this relationship well; *C. succinctus* which targets heather pollen, and *C. hederae* which is a specialist of ivy pollen (*Hedera helix*) (Figure 5, stress= 0.105). These bees display significantly different sterolomes despite being closely related (Figure 5A, PERMANOVA, F_1,31_=37.418, *p=*0.001, dispersion: F_1,31_=27.761, *p<*0.001), with species identity accounting for over 50% of the variation in the data (R^2^=0.547).

**Figure 5.**
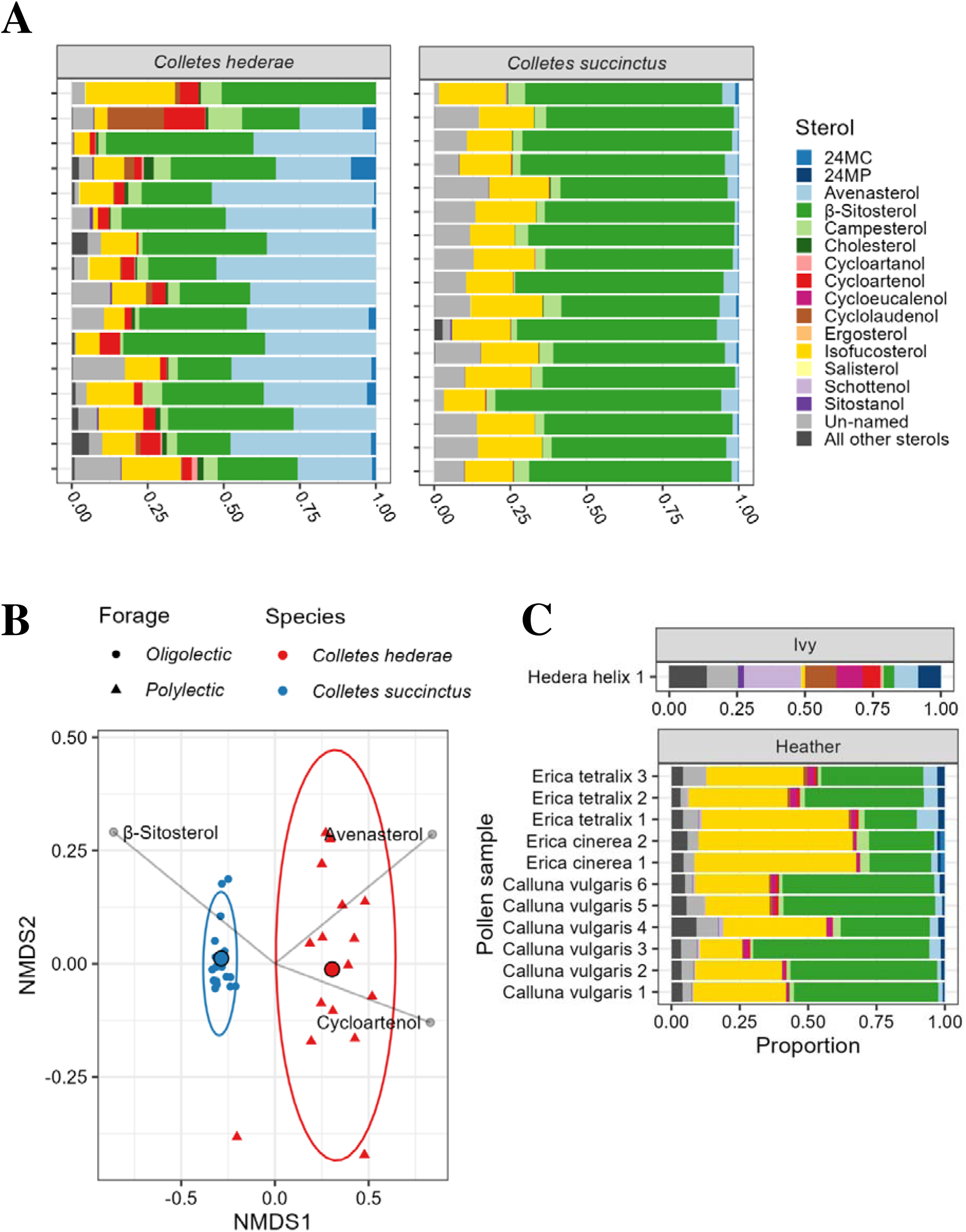
**A)** Sterol composition of all replicates of Colletes hederae and C. succinctus. Only major sterols are named with the remainder collapsed into single group. Colletes succinctus demonstrated much higher proportions of β-sitosterol whereas C. hederae contains higher proportions of avenasterol. Both species show a sterolome that is largely consistent across samples. **B)** NMDS of Colletes hederae and C. succinctus (stress= 0.105). Species show distinct sterolomes (PERMANOVA, F_1,31_=37.418, p=0.001) despite their relatedness. C. hederae is associated with higher avenasterol and cycloartenol whereas C. succinctus is associated with higher β-sitosterol. Colletes hederae shows a larger variation in sterolome, potentially due to its broader foraging habit. **C)** Sterol composition of the main pollen hosts of Colletes hederae and C. succinctus (Hedera helix and heather species). Only major sterols are named with the remainder collapsed into single group. The three Ericaceous heather species show a similar sterol profiles, dominated by isofucosterol and β-sitosterol. Hedera helix shows a much more variable sterol profile, however it is not the only forage plant of Colletes hederae.

The ivy bee, *Colletes hederae,* was most strongly associated with cycloartenol (IV=0.946, *p=*0.005) and avenasterol (IV= 0.856, *p=*0.005) which are both prominent sterols in ivy pollen (Baker et al., Biorxiv). In contrast, the profile of the heather colletes, *Colletes succinctus,* was strongly associated with β-sitosterol (IV= 0.814, *p=*0.005) and isofucostesterol (IV= 0.788, *p=*0.005) (full list of associations not shown). These associations reflect sterols which are at higher proportions in their respective pollens, indicating that diet strongly influences bee sterolome (Figure 5C).

### 3.4 Bumble bees selectively forage for pollen to maintain a consistent sterol profile

Bumble bees are generalist foragers and have been reported collecting pollen from a wide range of plants (Falk and Lewington, 2015). We compared the sterolome of corbiculate pollen collected by bumble bees to mean sterolomes of each bumble bee species (18) and to the sterolome of 295 different UK pollens (Baker et al. 2025). The sterol profile of corbiculate pollen was more similar to bumblebees than the hand collected pollens suggesting bumblebees target pollen specifically.

Corbiculate pollen and bumble bees showed higher 24-methylenecholesterol than hand collected pollen which had higher cycloartenol and which is either absent or a minor sterol in two bumble bees (Figure S3A, NMDS stress value= 0.179, PERMANOVA: F2,350=13.030, p=0.001). There was also a significant difference in the variance of the groups (F2,350=28.015, p <0.001; Note: as the pollen sample types covered a much wider taxonomic breadth than the bumblebees, this difference in variation is to be expected). Hand collected pollen was strongly associated with sterols containing a cyclopropane ring, cyclolaudenol and cycloartenol, whereas corbiculate pollen was strongly associated with the Δ5 sterols, desmosterol and cholesterol (Figure S3 B).

To understand whether bumblebees maintained the same sterol profile in their tissues under different floral landscapes, a set of widely distributed and common *Bombus* species was sampled from at least three floristically distinct sites. Significant intraspecies variation between sites would demonstrate plasticity in sterol profile, even in the absence of experimental conditions. Only the common carder bee, *Bombus pascuorum*, exhibited a significantly different sterol profile at different collection sites (NMDS stress = 0.177, PERMANOVA: F_3, 22_=1.818, *p<*0.050, dispersion: F_3,22_=0.504, *p>*0.500).

However, this was not strongly significant (*p=*0.046), and the variance explained by collection site was low (R^2^=0.199). Both the red-tailed bumblebee, *B. lapidarius,* and the buff-tailed bumblebee, *B. terrestris/lucorum,* showed no significant difference in sterolome between sites (*B. lapidarius*: NMDS stress = 0.123, PERMANOVA: F_2,13_=1.540, *p>*0.100, dispersion: F_2,13_=1.103, *p>*0.100. *B. terrestris/lucorum*: NMDS stress = 0.172, PERMANOVA: F_3, 35_=1.162, *p>*0.100, dispersion: F_3,35_=0.609, *p>*0.500). All NMDS biplots are shown in Figure S4. This suggests that bumblebees can adapt their foraging to fulfil sterol requirements in different landscapes.

The sterol requirements of bumblebees therefore appear to be consistent across species and different habitats. In addition, the sterol requirements of bumblebees appeared also consistent between sexes and body tagmata. Analysis of individual head, thorax and abdomens of bumblebees showed that the sterol profile of the head differed significantly from the abdomen (*p<*0.050) and thorax (*p<*0.010). However, the variance explained by both was very low and differences varied by species (Figure S5, NMDS: stress=0.150, PERMANOVA: Tagmata: F_2,181_=4.581, *p*=0.001, Species: F_5,181_=2.382, *p*=0.001, Variance explained: Tagmata: R^2^=0.045, Species: R^2^=0.059). *Bombus vestalis* had the most male/female specimens and there was no difference in sterolome between these groups (NMDS stress= 0.121, PERMANOVA: F_1,17_= 2.06, *p=*0.100, dispersion: F_1,17_=0.339, *p>*0.500). When all eight species with male and female specimens were compared there was also no difference between sexes. NMDS biplots are shown in Figure S6.

### 3.5 Bees are selective in their use of pollen sterols

While generalist bee species may be able to obtain optimum sterol ratios through foraging widely across species, specialist bee species are selective in their sterol use. The sterolomes of specialist bees analysed in this dataset do not directly resemble those in their pollens indicating they have mechanisms for selective uptake, potentially through sterol transporters in the midgut.

In support of this, the sterol profiles of four monolectic bees (*Melitta dimidiata*, *Macropis europaea*, *Melitta tricincta*, *Andrena florea*) and two species heavily dependent on a single pollen source (*Colletes hederae*, *C. halophilus*) were significantly different when compared to their pollen (F_1,56_=53.733, R^2^=0.231, *p=*0.001). There were also significant differences between bee-pollen pairing groups (F_5,56_=18.635, R^2^=0.401, *p=*0.001). The interaction between these effects was also significant (F_5,56_=5.854, R^2^=0.126, *p=*0.001), indicating that the relative difference between bee and its host pollen depended on the pairing of specialist and host (Figure S7 NMDS: stress = 0.162).

Furthermore, some of the specialist bee species contained sterols which were not detected in their pollen. For example, the bryony mining bee, *Andrena florea,* and the sainfoin bee, *Melitta dimidiata,* both contained ergosterol while their pollens did not (0.706% and 0.012% of total sterols respectively). The yellow-loosestrife bee, *Macropis europaea,* and the ivy bee, *Colletes hederae,* both contained cholesterol, which did not occur in their pollen (0.265% and 0.926% respectively) suggesting a capacity to convert sterols to cholesterol. Desmosterol was absent from ivy (*Hedera helix)* but present in the ivy bee, *Colletes hederae* (1.299%). Only the red bartsia bee, *Melitta tricincta,* contained sterols that were totally represented in its host species’ pollen, *Odontites vernus*.

### 3.6 Bees can adapt to the pollen sterols of different plants

The adaptation to using phytosterols in pollen has enabled bees to forgo the metabolic cost of sterol production. However, they appear limited in the phytsterols they are capable of using. Plants could, therefore, create ‘undesirable’ pollens to deter pollinivory by producing higher quantities of unusable sterols. It would, therefore, be beneficial to bees if they were capable of tolerating some of these ‘undesirable’ phytosterols in their tissues. Asteraceae was highlighted in Baker et al. *in prep* as producing lower proportions of Δ5 sterols and higher proportions of sterols with a B-ring cyclopropane substitution but still has specialist bee species associated with its pollen. To determine whether these bees appear to have adapted to these ‘undesirable’ sterols, we compared the sterolomes of generalist bee species, Asteraceae specialist bee species, and non-Asteraceae specialists to the pollen sterols of all three Asteraceae tribes (Cichorioideae, Asteroideae, and Carduoideae) (Figure 6A). Whilst bee and pollen sterolomes were distinct, the sterol profiles of Asteraceae specialist bees were most similar to the pollen of the Asteraceae, specifically Cichorioideae (PERMANOVA, stress=0.151; F_5,96_=16.041, *p=*0.001). Generalist bee species were significantly associated with 24-methylenecholesterol (ISA, Figure 6C). Non-Asteraceae specialist bees were significantly associated with β-sitosterol and Asteraceae specialists with cholesterol. Different Asteraceae tribe pollens were associated with at least one sterol with a B-ring cyclopropane substitution (CPR) each: Asteroideae with cycloartenol (IV=0.564, *p*=0.005), Carduoideae with cyclolaudenol (IV=0.568, *p=*0.005) and Cichorioideae with cycloartanol (IV=0.576, *p*=0.010). Carduoideae was associated with the most distinct pollen sterolome (Figure 6C).

**Figure 6.**
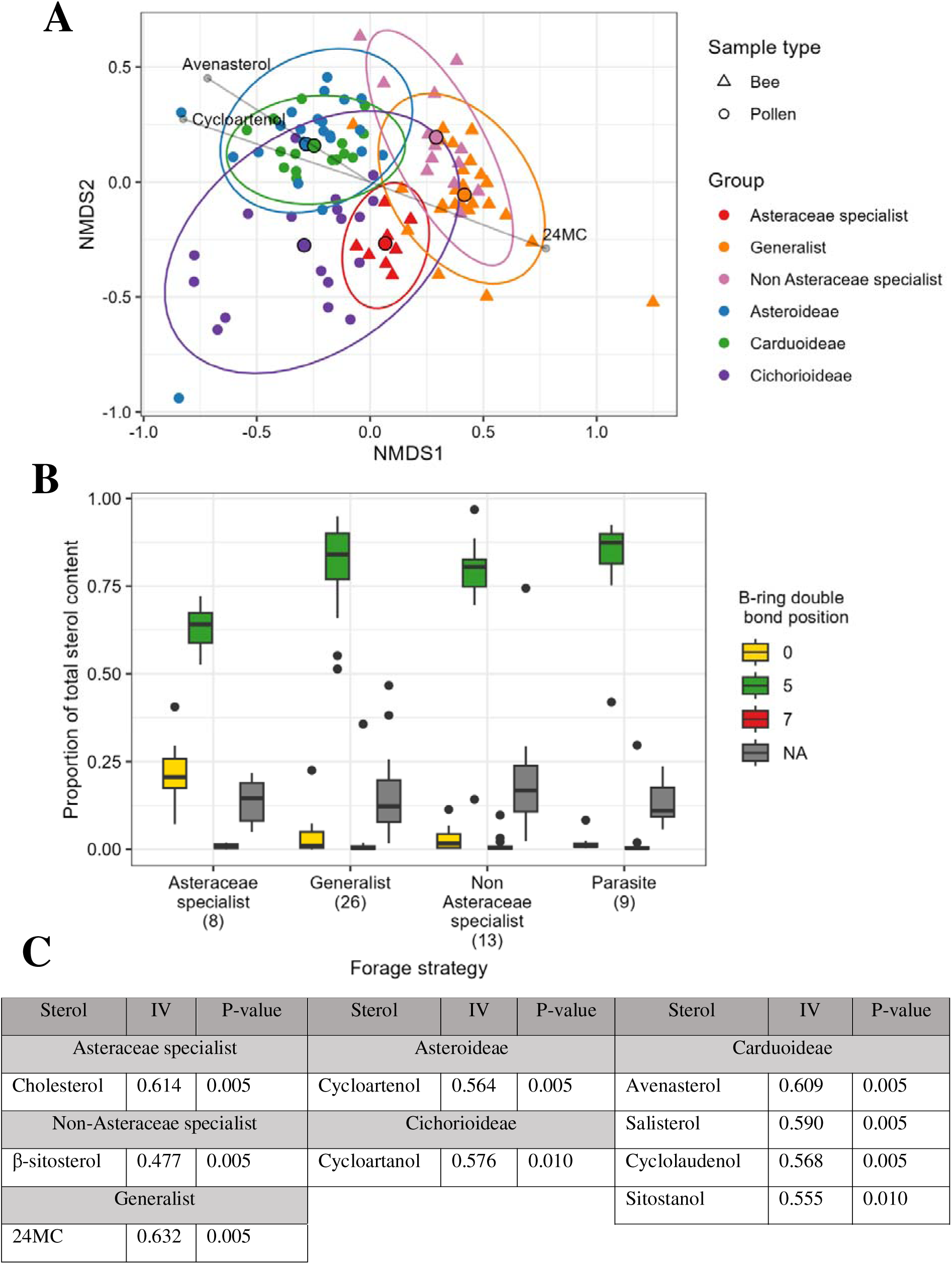
**A)** NMDS comparing the sterol profile of bees which specialise on Asteraceae pollen to those which do not (stress = 0.151). Bees and pollen species appear distinct in NMDS space. Most Asteraceae specialists target Cichorioideae and can be seen to cluster closer to these pollen than other Asteraceae. Groups were significantly different (PERMANOVA, stress=0.151; F_5,96_=16.041, p=0.001) and group identity explained nearly 50% of the variation in the dataset (R^2^=0.455). All pairwise comparisons between groups were significant (p<0.050) except Asteroideae versus Carduoideae pollen (p>0.100). Grey lines indicate significant (p<0.010) and strong correlations (absolute value >0.7). **B)** Proportion of sterols belonging to different double bond position groups across solitary bee species. Asteraceae specialists show a lower proportion of Δ5 sterols and higher proportion of sterols with a B-ring cyclopropane substitution (CPR) than the rest of the dataset. (CPR: H(3)=19.077, p<0.005, Δ5: H(3)=12.743, p<0.050, post-hoc Dunn tests: CPR (p<0.005), Δ5 (p<0.050)). No other comparisons were significantly different. **C)** Summary of Indicator Species Analysis results showing associations between groups in Figure 9A and individual sterols. All bee groups were associated with Δ5 sterol whereas each pollen group was associated with a sterol containing a cyclopropane ring.

When comparing all 56 bee species, Asteraceae specialists showed significantly lower proportions of Δ5 sterols and higher proportions of sterols with a B-ring cyclopropane substitution (CPR) than non-Asteraceae specialist, generalist and parasitic bee species (CPR: H(3)=19.077, *p<*0.005, Δ5: H(3)=12.743, *p<*0.050, *post-hoc* Dunn tests: CPR (*p<*0.005), Δ5 (*p<*0.050)) (Figure 6B).

## 4 DISCUSSION

This study represents the largest survey of wild bee sterolomes to date. The whole-body tissues of all 78 bee species analysed were dominated by phytosterols rather than cholesterol, indicating that in general, bees do not convert dietary sterols into cholesterol. Instead, they substitute phytosterols into their cell membranes. Generalist and specialist species alike share a sterolome dominated by 24-methylenecholesterol, β-sitosterol and isofucosterol, as predicted by previous work (Svoboda and Lusby, 1986; Vanderplanck, Zerck, *et al*., 2020). The 18 separately analysed *Bombus* species showed a consistent sterolome comparable to those reported in honey bees (Svoboda *et al*., 1980). These sterols are available in high proportions across a wide taxonomic range of floral pollen. However, sterol abundance alone does not lead to their use by bees; cycloartenol, cycloeucalenol and obtusifoliol were abundant across a range of plants but largely absent in bees. Our data implies that these three sterols are therefore best suited to bee physiological requirements, perhaps in part due to their Δ5 B-ring double bond. The lack of phylogenetic signal in the dataset indicates the bee sterolome is driven by other forces such as foraging choice and life history, creating significant differences between event closely related bees.

Sterols are a stabilizing compound in cell membranes which impact membrane permeability and fluidity (Reviewed in Dufourc, 2008). The phytosterols stigmasterol and β-sitosterol can form rafts and maintain a stable membrane fluidity across a wider temperature range than cholesterol. This is likely an adaptation to the wider range of temperature conditions plants experience compared to animals. Therefore, using these longer chain phytosterols in place of cholesterol could make also bees more tolerant of cold conditions (Knittelfelder *et al*., 2020). In addition to the three sterols mentioned above, several other sterols, campesterol, cycloartanol, avenasterol, were present in proportions (>30%) suggesting their incorporation into membranes. Cycloartenol, has been shown to substitute for cholesterol when maintaining membrane dynamics in model tissues but was not dominant in bee tissues (Dufourc, 2008).

However, insects also require sterols for the synthesis of steroid hormones: in most species, cholesterol is converted into hormones (Jing and Behmer, 2020). Honeybees need cholesterol and campesterol to synthesize the hormones 20-hydroxyecdysone and makisterone A (Feldlaufer *et al*., 1986). Other dietary sterols can also be essential to successful development in bees, as seen with the reliance of a stingless bee on ergosterol to pupate (Paludo *et al*., 2018). The few bee species that have been studied for their steroid hormone precursors show that many bees may share the trait of using campesterol as the basis for production of the alternative moulting hormone, makisterone A (Feldlaufer *et al*., 1993; Paludo *et al*., 2018). In our data, the phylogenetic trend seen in campesterol, where closely related species showed similar proportions, may therefore be due to its importance in hormone production. Further, the absence of cholesterol or campesterol from some species (*Andrena wilkella*, *Nomada flavopicta*, *Halictus rubicundus*, *Lasioglossum smeathmanellum*, *Coelioxys inermis* and *Ceratina cyanea*.) could indicate that these species use other moulting hormone synthesis pathways (Feldlaufer *et al*., 1986; Furse, Koch, *et al*., 2023). It is however important to note that these species were all represented by a single sample. Therefore, it is possible cholesterol and campesterol were too low to be detectable in these samples.

It was shown that collection site, and therefore available plant community, did not affect sterol profile in two out of three bumble bee species analysed. This suggests bumble bees will maintain an optimum sterol profile through foraging on different pollen, demonstrating that even generalists are selective in their sterol usage. The seemingly stable ratio between sites may therefore represent a compromise between readily available dietary sterols and a physiological optimum (Knittelfelder *et al*., 2020).

This strategy may contribute to the ecological success of bumble bees, allowing them to forage in an extremely wide range of habitats, using pollen from many different plant species. Within the bumble bee dataset, the interspecies differences and species-sterol associations were the result of small variations in proportions of different sterols, likely a result of their similar foraging and life history strategies. As such, the ecological relevance of these differences may be limited.

Corbiculate pollen collected from bumble bees showed a profile distinct from other hand-collected pollens, correlating with higher isofucosterol and 24-methylenecholesterol. It is therefore possible that bees are targeting certain sterols in pollen, though there is currently no evidence to suggest they can detect sterols directly. There are also other potential causes of variation between hand and bee collected pollens; bumble bees can change their pollen foraging target during a single trip (Martínez-Bauer *et al*., 2021) and they add saliva to collected pollen to aid compaction. Further, the bees analysed in this study will likely have collected different pollen to those they were raised on.

Despite shared similarities, the sterolomes of generalist and specialist bee species were distinct. For instance, polylectic *Andrena* species demonstrated higher proportions of 24-methylenecholesterol than oligo/monolectic species. The generalist’s ability to forage across a wide range of flowers could facilitate greater access to desirable sterols such as 24-methylenecholesterol, but at a cost of competition for resources. In contrast, specialisation would reduce competition but achieve a less ideal ratio of sterols. The ability of specialist bees to adapt their sterolome to make best use of their pollen host, as seen in *Colletes*, suggests there is plasticity in sterol usage in bees, operating within physiological constraints.

This is made clear in the monolectic and highly specialised bees which showed sterolomes distinct from their host pollen, indicating that bees have mechanisms for the selective uptake of sterols from diet. Some bee species contained sterols absent from their pollen, indicating that the specialist plant-bee relationship is flexible and bees are using pollen from additional plant species. It could alternatively indicate that a failed detection of these compounds in our samples. In addition, the sterol requirements of specialist bees could be fulfilled by plants other than their chosen species; for this reason, we expect that the sterols are not what are driving specialization in host-pollinator relationships. In fact, relying on a single pollen host, a strategy which makes them highly vulnerable to changes in flowering times and distribution, could be risky for any species of bee. For example, *Macropis europaea* also collects fatty floral oils from its main pollen plant *Lysimachia vulgaris* to waterproof its nests in damp soils (Falk and Lewington, 2015).

We were able to identify potential specialist sterol adaptation most distinctly in Asteraceae specialist bees due to their frequency in the dataset. The pollen sterols of Asteraceae flowers display lower levels of Δ5 sterols such as 24-methylenecholesterol, β-sitosterol and isofucosterol and higher levels of sterols with Δ8 B-ring bonds and cyclopropane rings (Baker et al. *in prep*). This may be a potential cause of their less frequent usage by generalist bee species (Müller and Kuhlmann, 2008; Vanderplanck, Gilles, *et al*., 2020) as these sterols do not appear suitable for the majority of bee species and may prevent successful growth and development (Jing, Grebenok and Behmer, 2012; Behmer, 2017). Our results showed that this trend was mirrored in the sterolomes of Asteraceae specialist bees which often had higher levels of cycloartanol and cholesterol and lower proportions Δ5 sterols. The ability to incorporate much higher levels of cycloartanol in their tissues than most other bee species suggests these species have adapted to Asteraceae pollen. However, these bees still maintained a dominance of Δ5 sterol indicating there are limits to sterolome adaptation in bees. The ability to overcome the nutritional deficiencies of Asteraceae pollen may have allowed these bees to benefit from the widespread and abundant availability of these flowers.

Cichorioideae flowers produced some of the highest cholesterol proportions presented in Baker et al. *in prep* (*Helminthotheca echioides*: 60.05%, *Crepis capillaris*: 44.21%, *Leontodon saxatilis*: 31.33%, *Crepis vesicaria*: 27.77%). This has been proposed as facilitating the transition from carnivory to pollinivory in Apidae evolution (Santerre, 2023). As bees evolved from carnivorous ancestors to pollinivorous diets, cholesterol rich pollens may have provided an intermediary step in the transition to pollinivory, before evolving to use longer chain phytosterols in their tissues, as suggested by Dötterl and Vereecken (2010). Dasypoda species, which occupy a basal position in bee evolution, are also ancestrally oligolectic on Asteraceae flowers (Michez *et al*., 2008).

The results of this analysis do not reveal a singular clear trend in the sterol profile of wild bees. There is no evidence for a strong phylogenetic signal in most sterols, therefore, the drivers of sterol profile in bees would appear to be ecologically, rather than phylogenetically, constrained but with some plasticity. However, there are multiple instances of sterol profiles differing from those expected by ecological drivers, for instance where monolectic bee species sterol profiles do not match the sterol proportions of their pollen host. It may be that selective uptake and sequestration means proportional differences are present between pollen and bee sterol profiles and there are other nutritional or ecological factors which strongly influence pollen foraging choices. Future work can build on these data by experimentally manipulating the diets of bees to determine if they are fixed to a profile of specific sterols or if they demonstrate greater flexibility. The ecological and economic importance of wild bees makes further understanding their nutritional requirements an important step understanding the impact of floral community change on bee populations.

