## Supplementary Tables and Figures for "Specialization and adaptation in pollen sterol use by wild bees"

**Table S1.** Median, interquartile range, minimum and maximum percentages for all 19 sterols detected, including six unidentified sterols. Isofucosterol, β-sitosterol and 24-methylenecholesterol show the highest median values and are the only sterols common to all species. Data was calculated from mean species proportions.

| **Sterol** | **Median (%)** | **IQR** | **Max (%)** | **Min (%)** | **No. of Species detected in** |
| --- | --- | --- | --- | --- | --- |
| Isofucosterol | 21.797 | 13.293 | 38.151 | 0.164 | 56 |
| β-Sitosterol | 20.581 | 19.168 | 74.291 | 2.307 | 56 |
| 24MC | 12.633 | 36.718 | 88.009 | 0.378 | 56 |
| ST(29:2)A | 7.928 | 8.575 | 23.566 | 0.000 | 54 |
| Campesterol | 4.802 | 5.029 | 31.174 | 0.000 | 54 |
| ST(30:1)A | 1.255 | 0.876 | 4.295 | 0.000 | 55 |
| ST(28:1)B | 0.639 | 1.456 | 43.893 | 0.000 | 53 |
| ST(29:1)A | 0.541 | 0.756 | 4.502 | 0.000 | 50 |
| Avenasterol | 0.382 | 0.977 | 35.653 | 0.000 | 49 |
| Desmosterol | 0.382 | 0.751 | 6.201 | 0.000 | 53 |
| Cholesterol | 0.312 | 1.717 | 20.012 | 0.000 | 51 |
| Cycloartenol | 0.252 | 0.513 | 4.221 | 0.000 | 46 |
| Cycloartanol | 0.218 | 3.193 | 38.288 | 0.000 | 43 |
| ST(28:1)A | 0.212 | 0.431 | 2.773 | 0.000 | 46 |
| Sitostanol | 0.175 | 0.208 | 3.845 | 0.000 | 51 |
| ST(30:1)B | 0.165 | 0.511 | 51.694 | 0.000 | 49 |
| Cyclolaudenol | 0.145 | 0.294 | 6.344 | 0.000 | 49 |
| Ergosterol | 0.027 | 0.072 | 12.697 | 0.000 | 39 |
| Salisterol | 0.000 | 0.022 | 1.719 | 0.000 | 26 |

**Table S2.** Median, inter-quartile range (IQR), maximum and minimum of all sterols detected in whole Bombus samples. Any sterols not detected in both runs were removed Number of species each sterol was found in also listed (maximum 18). Arranged by median value. Un-named sterols are labelled with carbon chain length (CC) and double bond count (DB) as (CC:DB).

| **Sterol** | **Median** | **IQR** | **Max** | **Min** | **Species detected in** |
| --- | --- | --- | --- | --- | --- |
| 24MC | 34.898 | 14.758 | 58.509 | 8.796 | 18 |
| Isofucosterol | 27.568 | 7.472 | 34.871 | 16.842 | 18 |
| β-Sitosterol | 22.619 | 7.370 | 49.817 | 6.673 | 18 |
| Campesterol | 6.644 | 2.709 | 18.139 | 0.918 | 18 |
| Ergosterol | 2.099 | 7.603 | 19.000 | 0.033 | 18 |
| Desmosterol | 1.178 | 0.904 | 5.386 | 0.092 | 18 |
| Cycloartenol | 0.501 | 0.245 | 0.979 | 0.021 | 18 |
| Cholesterol | 0.353 | 0.359 | 1.253 | 0.038 | 18 |
| Cyclolaudenol | 0.139 | 0.127 | 0.881 | 0.015 | 18 |

**Table S3.** Summary of the number of sterols detected in each bee species and the sterols which were missing if they contained less than the maximum of 19.

| **Species** | **No. of sterols** | **Missing sterols** |
| --- | --- | --- |
| *Andrena bicolor* | 19 | None |
| *Andrena cineraria* | 19 | None |
| *Andrena fuscipes* | 19 | None |
| *Andrena hattorfiana* | 19 | None |
| *Anthophora plumipes* | 19 | None |
| *Bombus sylvarum* | 19 | None |
| *Bombus terrestris* | 19 | None |
| *Colletes halophilus* | 19 | None |
| *Colletes hederae* | 19 | None |
| *Dasypoda hirtipes* | 19 | None |
| *Epeolus cruciger* | 19 | None |
| *Lasioglossum brevicorne* | 19 | None |
| *Macropis europaea* | 19 | None |
| *Melitta haemorrhoidalis* | 19 | None |
| *Melitta tricincta* | 19 | None |
| *Nomada fucata* | 19 | None |
| *Osmia leaiana* | 19 | None |
| *Andrena flavipes* | 18 | Salisterol |
| *Andrena florea* | 18 | Avenasterol |
| *Andrena fulvago* | 18 | Salisterol |
| *Anthophora furcata* | 18 | ST(28:1)A |
| *Anthophora quadrimaculata* | 18 | Salisterol |
| *Bombus pascuorum* | 18 | Salisterol |
| *Bombus vestalis* | 18 | ST(30:1)B |
| *Chelostoma florisomne* | 18 | Salisterol |
| *Colletes succinctus* | 18 | Ergosterol |
| *Melitta dimidiata* | 18 | Cycloartanol |
| *Melitta leporina* | 18 | Salisterol |
| *Osmia bicornis* | 18 | Salisterol |
| *Andrena fulva* | 17 | Cycloartanol, salisterol |
| *Andrena nitida* | 17 | Cycloartenol, salisterol |
| *Andrena scotica* | 17 | Cycloartenol, salisterol |
| *Bombus lucorum* | 17 | Cycloartanol, alisterol |
| *Epeolus variegatus* | 17 | Salisterol, ST(29:1)A |
| *Halictus tumulorum* | 17 | Ergosterol, salisterol |
| *Megachile willughbiella* | 17 | Cycloartanol, salisterol |
| *Melecta albifrons* | 17 | Cycloartanol, salisterol |
| *Nomada rufipes* | 17 | Ergosterol, Salisterol |
| *Andrena denticulata* | 16 | Ergosterol, salisterol, ST(28:1)B |
| *Andrena humilis* | 16 | Salisterol, sitostanol, ST(28:1)B |
| *Colletes daviesanus* | 16 | Ergosterol, salisterol, ST(28:1)A |
| *Hylaeus communis* | 16 | Cyclolaudenol, ergosterol, salisterol, |
| *Lasioglossum villosulum* | 16 | Cycloartenol, salisterol, ST(28:1)A |
| *Sphecodes monilicornis* | 16 | Cycloartenol, ergosterol, salisterol |
| *Bombus hortorum* | 15 | Cycloartanol, cyclolaudenol, ergosterol, salisterol |
| *Bombus pratorum* | 15 | Cycloartanol, cycloartenol, ergosterol, salisterol |
| *Chelostoma campanularum* | 15 | Ergosterol, salisterol, ST(28:1)A, ST(29:1)A |
| *Heriades truncorum* | 15 | Ergosterol, sitostanol, ST(28:1)A, ST(29:1)A |
| *Coelioxys inermis* | 14 | Avenasterol, cholesterol, cycloartenol, ergosterol, ST(30:1)B |
| *Hylaeus signatus* | 14 | Cycloartanol, cycloartenol, ergosterol, ST(28:1)A, ST(30:1)B |
| *Anthidium manicatum* | 13 | Cycloartanol, cyclolaudenol, salisterol, sitostanol, ST(28:1)A, ST(30:1)B |
| *Andrena wilkella* | 12 | Avenasterol, Cholesterol, Cycloartanol, Desmosterol, Ergosterol, Salisterol, ST(28:1)A |
| *Lasioglossum smeathmanellum* | 10 | Avenasterol, cholesterol, cycloartanol, cyclolaudenol, ergosterol, salisterol, sitostanol, ST(29:1)A, ST(30:1)B, |
| *Halictus rubicundus* | 9 | Avenasterol, campesterol, cholesterol, cycloartanol, cycloartenol, cyclolaudenol, desmosterol, salisterol, sitostanol, ST(29:2)A |
| *Ceratina cyanea* | 8 | Avenasterol, campesterol, cycloartenol, cyclolaudenol, ergosterol, ST(28:1)A, ST(28:1)B, ST(29:1)A, ST(29:2)A, ST(30:1)A, ST(30:1)B |
| *Nomada flavopicta* | 8 | Avenasterol, cholesterol, cycloartanol, cycloartenol, cyclolaudenol, desmosterol, ergosterol, salisterol, ST(28:1)A, ST(29:1)A, ST(30:1)B |

**Table S4.** Phylogenetic signal analysis results for wild bees dataset. Only three sterols showed phylogenetic signal for both metrics (Blomberg’s K and Pagels λ). All significant values are highlighted in grey.

| **Group** | **K** | **λ** | **P K** | **P λ** |
| --- | --- | --- | --- | --- |
| Desmosterol | 0.327 | 0.790 | 0.044 | 0.002 |
| Campesterol | 0.525 | 1.012 | 0.001 | 0.001 |
| ST(28:1)B | 0.479 | 0.758 | 0.045 | 0.001 |
| ST(28:1)A | 0.310 | 0.535 | 0.047 | 0.192 |
| 24MC | 0.288 | 0.178 | 0.015 | 0.363 |
| Sitostanol | 0.397 | 0.997 | 0.079 | 0.038 |
| Salisterol | 0.628 | 1.007 | 0.066 | 0.001 |
| 28 (carbon chain length) | 0.289 | 0.290 | 0.010 | 0.123 |
| Δ0 (B-ring saturation) | 0.397 | 0.997 | 0.094 | 0.038 |

***Table S5.*** *Retention time, mass to charge ratio (m/z) and structure description of sterols (carbon count (CC): double bond equivalents (DB)) used in Quality Assurance (QA) mixture.*

| **Sterol** | **Structure** | **m/z** | **Retention time (min)** |
| --- | --- | --- | --- |
| 24-methylenecholesterol | (28:2) | 398.3549 | 5.9 |
| 24-methylenecycloartenol | (31:2) | 440.4018 | 7.5 |
| Anthelsterol | (29:3) | 410.3549 | 5.3 |
| Avenasterol | (29:2) | 412.3705 | 6.9 |
| β-sitosterol | (29:1) | 414.3862 | 9.7 |
| Brassicasterol | (28:2) | 398.3549 | 6.6 |
| Campesterol | (28:1) | 400.3705 | 8.2 |
| Cholesterol | (27:1) | 386.3549 | 7.0 |
| Cycloartenol | (30:2) | 426.3862 | 8.3 |
| Cycloeucalenol | (30:2) | 426.3862 | 7.7 |
| Cyclolaudenol | (31:2) | 440.4018 | 9.5 |
| Desmosterol | (27:2) | 384.3392 | 5.1 |
| Episterol | (28:2) | 398.3549 | 5.7 |
| Ergosterol | (28:3) | 396.3392 | 5.5 |
| Isofucosterol | (29:2) | 412.3705 | 7.5 |
| Schottenol | (29:1) | 414.3862 | 9.0 |
| Sitostanol | (29:0) | 416.4018 | 11.1 |
| Spinasterol | (29:2) | 412.3705 | 8.0 |
| Stigmasterol | (29:2) | 412.3705 | 8.5 |

| Family | Subfamily | Tribe | Genus | Species |
| --- | --- | --- | --- | --- |
| Andrenidae | Andreninae | Andrenini | *Andrena* | *Andrena bicolor* |
| Andrenidae | Andreninae | Andrenini | *Andrena* | *Andrena cineraria* |
| Andrenidae | Andreninae | Andrenini | *Andrena* | *Andrena denticulata* |
| Andrenidae | Andreninae | Andrenini | *Andrena* | *Andrena flavipes* |
| Andrenidae | Andreninae | Andrenini | *Andrena* | *Andrena florea* |
| Andrenidae | Andreninae | Andrenini | *Andrena* | *Andrena fulva* |
| Andrenidae | Andreninae | Andrenini | *Andrena* | *Andrena fulvago* |
| Andrenidae | Andreninae | Andrenini | *Andrena* | *Andrena fuscipes* |
| Andrenidae | Andreninae | Andrenini | *Andrena* | *Andrena hattorfiana* |
| Andrenidae | Andreninae | Andrenini | *Andrena* | *Andrena humilis* |
| Andrenidae | Andreninae | Andrenini | *Andrena* | *Andrena nitida* |
| Andrenidae | Andreninae | Andrenini | *Andrena* | *Andrena scotica* |
| Andrenidae | Andreninae | Andrenini | *Andrena* | *Andrena wilkella* |
| Megachilidae | Megachilinae | Anthidiini | *Anthidium* | *Anthidium manicatum* |
| Apidae | Anthophorinae | na | *Anthophora* | *Anthophora furcata* |
| Apidae | Anthophorinae | na | *Anthophora* | *Anthophora plumipes* |
| Apidae | Anthophorinae | na | *Anthophora* | *Anthophora quadrimaculata* |
| Apidae | Apinae | Bombini | *Bombus* | *Bombus hortorum* |
| Apidae | Apinae | Bombini | *Bombus* | *Bombus lucorum* |
| Apidae | Apinae | Bombini | *Bombus* | *Bombus pascuorum* |
| Apidae | Apinae | Bombini | *Bombus* | *Bombus pratorum* |
| Apidae | Apinae | Bombini | *Bombus* | *Bombus sylvarum* |
| Apidae | Apinae | Bombini | *Bombus* | *Bombus terrestris* |
| Apidae | Apinae | Bombini | *Bombus* | *Bombus vestalis* |
| Apidae | Xylocopinae | Ceratinini | *Ceratina* | *Ceratina cyanea* |
| Megachilidae | Megachilinae | Osmiini | *Chelostoma* | *Chelostoma campanularum* |
| Megachilidae | Megachilinae | Osmiini | *Chelostoma* | *Chelostoma florisomne* |
| Megachilidae | Megachilinae | Megachilini | *Coelioxys* | *Coelioxys inermis* |
| Colletidae | Colletinae | na | *Colletes* | *Colletes daviesanus* |
| Colletidae | Colletinae | na | *Colletes* | *Colletes halophilus* |
| Colletidae | Colletinae | na | *Colletes* | *Colletes hederae* |
| Colletidae | Colletinae | na | *Colletes* | *Colletes succinctus* |
| Melittidae | Dasypodainae | Dasypodaini | *Dasypoda* | *Dasypoda hirtipes* |
| Apidae | Nomadinae | Epeolini | *Epeolus* | *Epeolus cruciger* |
| Apidae | Nomadinae | Epeolini | *Epeolus* | *Epeolus variegatus* |
| Halictidae | Halictinae | Halictini | *Halictus* | *Halictus rubicundus* |
| Halictidae | Halictinae | Halictini | *Halictus* | *Halictus tumulorum* |
| Megachilidae | Megachilinae | Osmiini | *Heriades* | *Heriades truncorum* |
| Colletidae | Hylaeinae | na | *Hylaeus* | *Hylaeus communis* |
| Colletidae | Hylaeinae | na | *Hylaeus* | *Hylaeus signatus* |
| Halictidae | Halictinae | Halictini | *Lasioglossum* | *Lasioglossum brevicorne* |
| Halictidae | Halictinae | Halictini | *Lasioglossum* | *Lasioglossum smeathmanellum* |
| Halictidae | Halictinae | Halictini | *Lasioglossum* | *Lasioglossum villosulum* |
| Melittidae | Melittinae | Macropidini | *Macropis* | *Macropis europaea* |
| Megachilidae | Megachilinae | Megachilini | *Megachile* | *Megachile willughbiella* |
| Apidae | Nomadinae | Melectini | *Melecta* | *Melecta albifrons* |
| Melittidae | Melittinae | Melittini | *Melitta* | *Melitta dimidiata* |
| Melittidae | Melittinae | Melittini | *Melitta* | *Melitta haemorrhoidalis* |
| Melittidae | Melittinae | Melittini | *Melitta* | *Melitta leporina* |
| Melittidae | Melittinae | Melittini | *Melitta* | *Melitta tricincta* |
| Apidae | Nomadinae | Nomadini | *Nomada* | *Nomada flavopicta* |
| Apidae | Nomadinae | Nomadini | *Nomada* | *Nomada fucata* |
| Apidae | Nomadinae | Nomadini | *Nomada* | *Nomada rufipes* |
| Megachilidae | Megachilinae | Osmiini | *Osmia* | *Osmia bicornis* |
| Megachilidae | Megachilinae | Osmiini | *Osmia* | *Osmia leaiana* |
| Halictidae | Halictinae | Sphecodini | *Sphecodes* | *Sphecodes monilicornis* |

***Table S6.*** *Taxonomic information for all 56 bee species analysed, taken from Henríquez-Piskulich, Hugall and Stuart-Fox (2024) supplementary.*

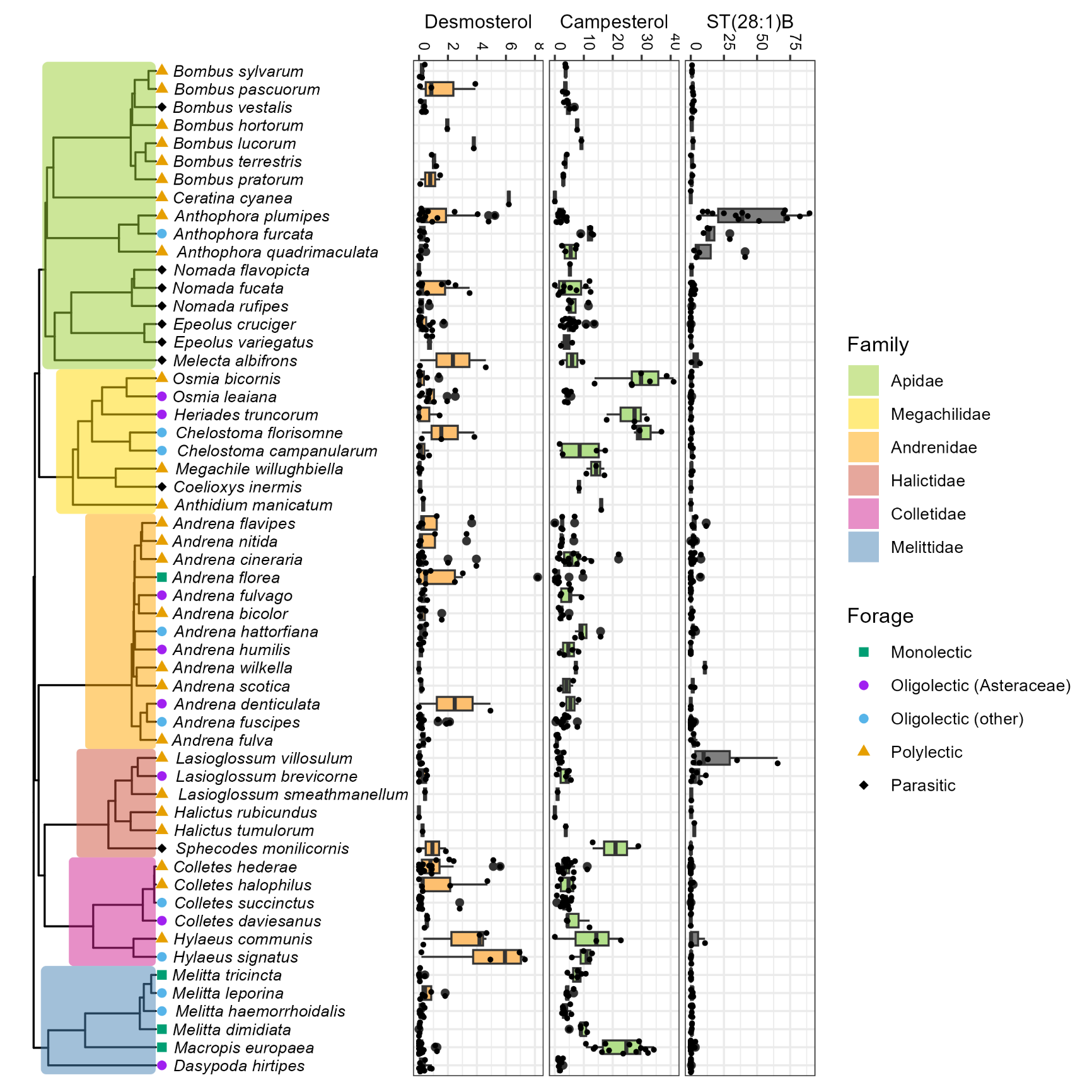

**Figure S1.** Phylogeny showing the proportion of three sterols shown to have phylogenetic signal across the 56 wild bee species analysed. Campesterol is higher in some Megachilidae species and lower in Apidae and most Halictidae. Whereas the distribution of higher desmosterol appears less conserved.

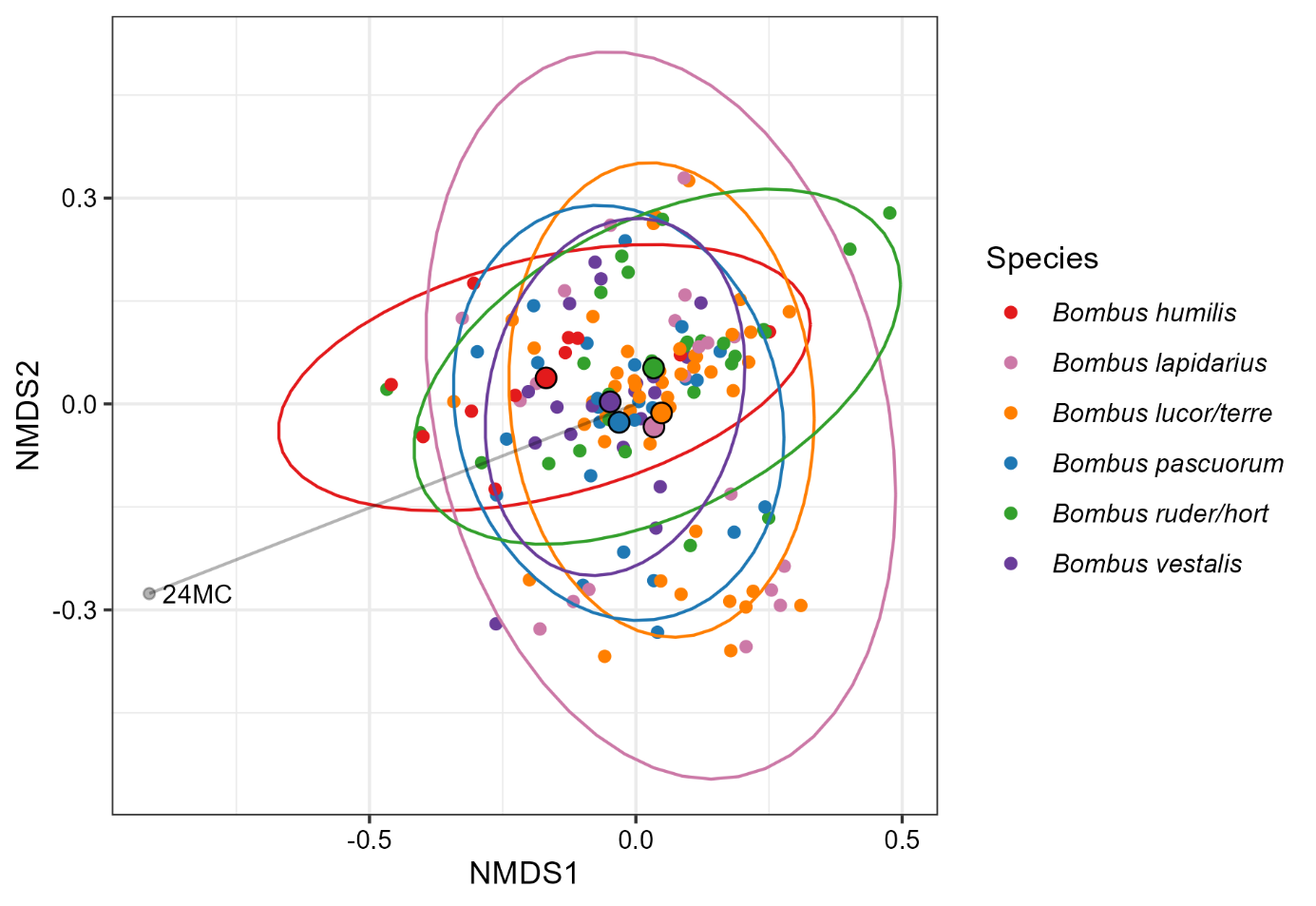

***Figure S2.*** *First two dimensions of a 3D NMDS carried out using the six most replicated* Bombus *species in the dataset (stress =0.128). Subsequent analysis showed significant inter-species differences.*

**Figure S3**

***A)*** *Comparison of whole-body bumble bee sterols with corbiculate and hand collected pollens. Bumble bee tissue sterols show a composition shared by a small proportion of hand collected pollens but a more similar mean profile to corbiculate pollen. Both bumble bees and corbiculate pollen showed higher levels of 24-methylenecholesterol than hand collected pollens. Both pollen groups displayed much larger variation in sterolome than the bumble bee specimens, likely a result of their wider taxonomic breadth. All groups were significantly different from each other (PERMANOVA: F_2,350_=13.030,* p*=0.001). Grey lines show strong (>0.7 absolute value) and significant (p<0.010) correlations. NMDS stress value = 0.179.*

***B)*** *Summary of Indicator Species Analysis results showing associations between groups in Figure 6A and individual sterols. Hand collected pollen was associated with longer chain (C30-31) sterols containing a cyclopropane ring in the B-ring of the steroid nucleus. In contrast corbiculate pollen was associated with C27 Δ5 sterols.*

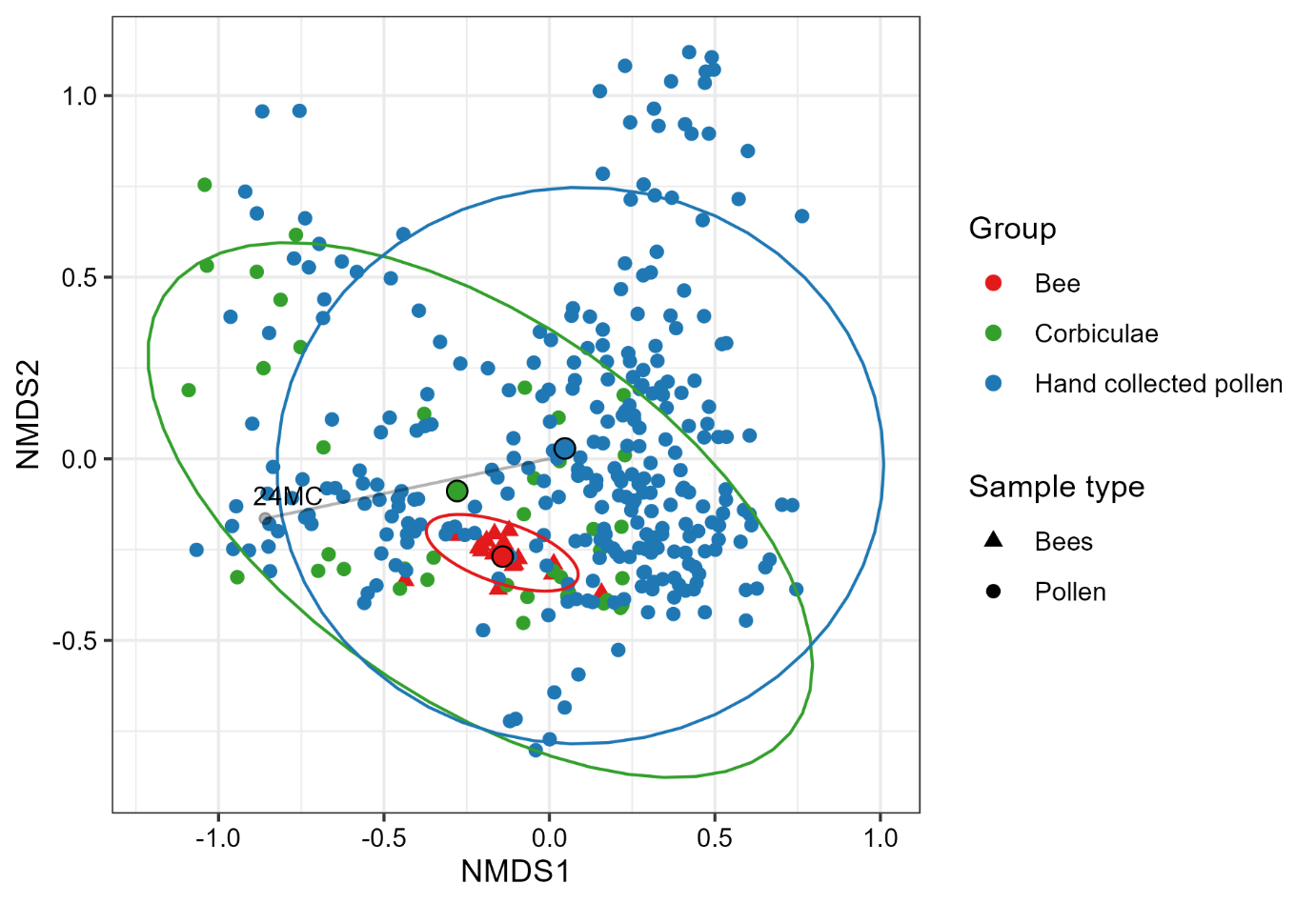

**A**

**B**

| Sterol | IV | P-value | Sterol | IV | P-value | Sterol | IV | P-value |
| --- | --- | --- | --- | --- | --- | --- | --- | --- |
| *Bumble bee whole bodies* | | | *Corbiculate pollen* | | | *Hand collected pollen* | | |
| Ergosterol | 0.795 | 0.005 | Desmosterol | 0.753 | 0.020 | Cyclolaudenol | 0.835 | 0.005 |
|  |  |  | Cholesterol | 0.730 | 0.005 | Cycloartenol | 0.779 | 0.005 |

**Figure S4:** *NMDS of the bumblebee species collected from multiple UK counties, coloured by collection county and annotated with sterols which showed strong (>0.7 absolute value) and significant (*p*<0.010) associations. All three species showed high overlap in sterolome between collection sites;* Bombus pascuorum *showed weak significant differences in sterol profile among sites (NMDS stress = 0.177, PERMANOVA: F_3, 22_=1.818,* p*<0.050). However,* Bombus lapidarius *and* B. terrestris/lucorum *did not show a difference in sterolome between sites (*B. lapidarius*: NMDS stress = 0.123, PERMANOVA: F_2,13_=1.540,* p*>0.100.* B. terrestris/lucorum*: NMDS stress = 0.172, PERMANOVA: F_3, 35_=1.162,* p*>0.100).*

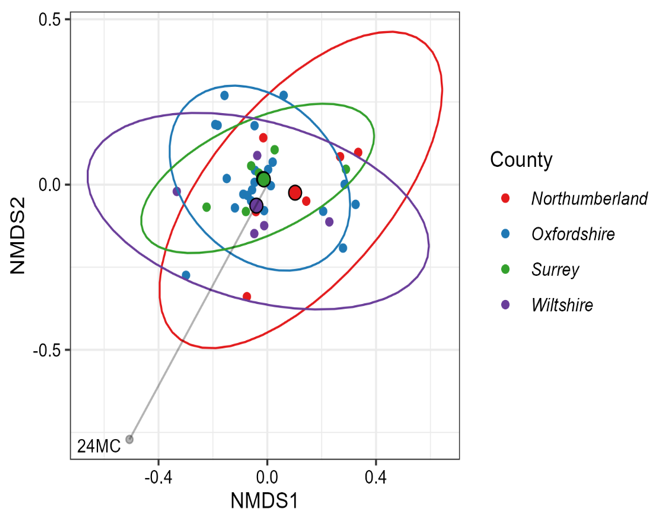

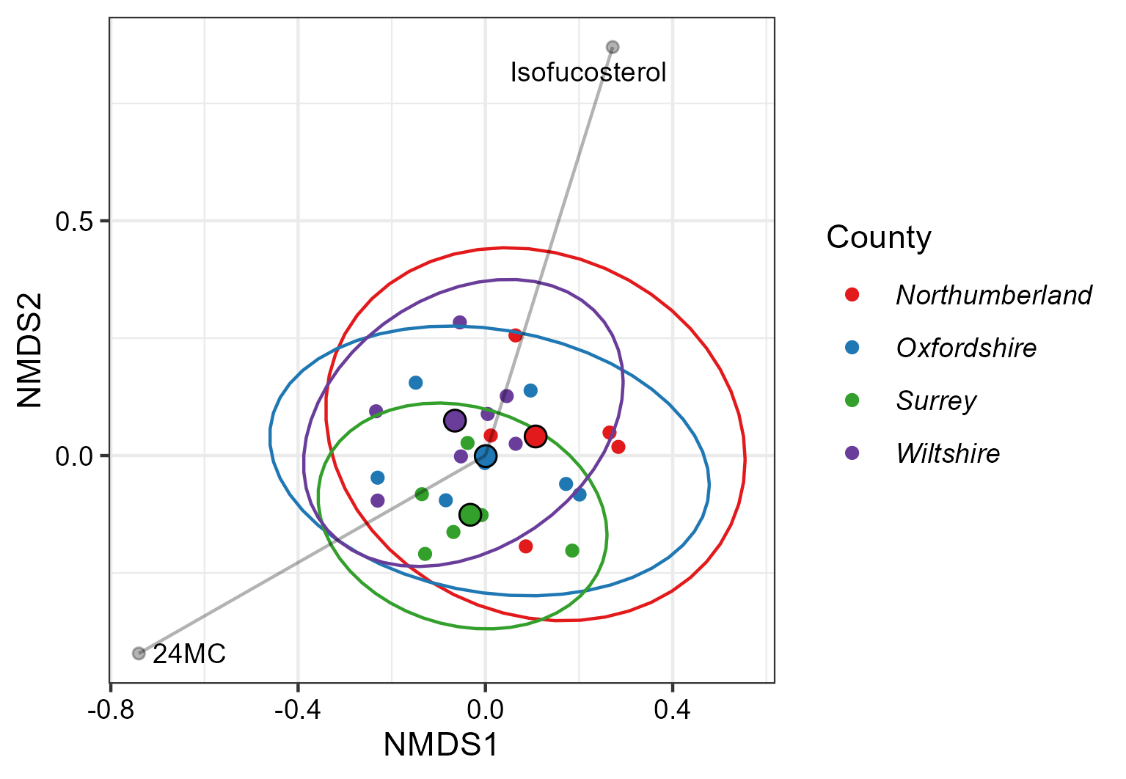

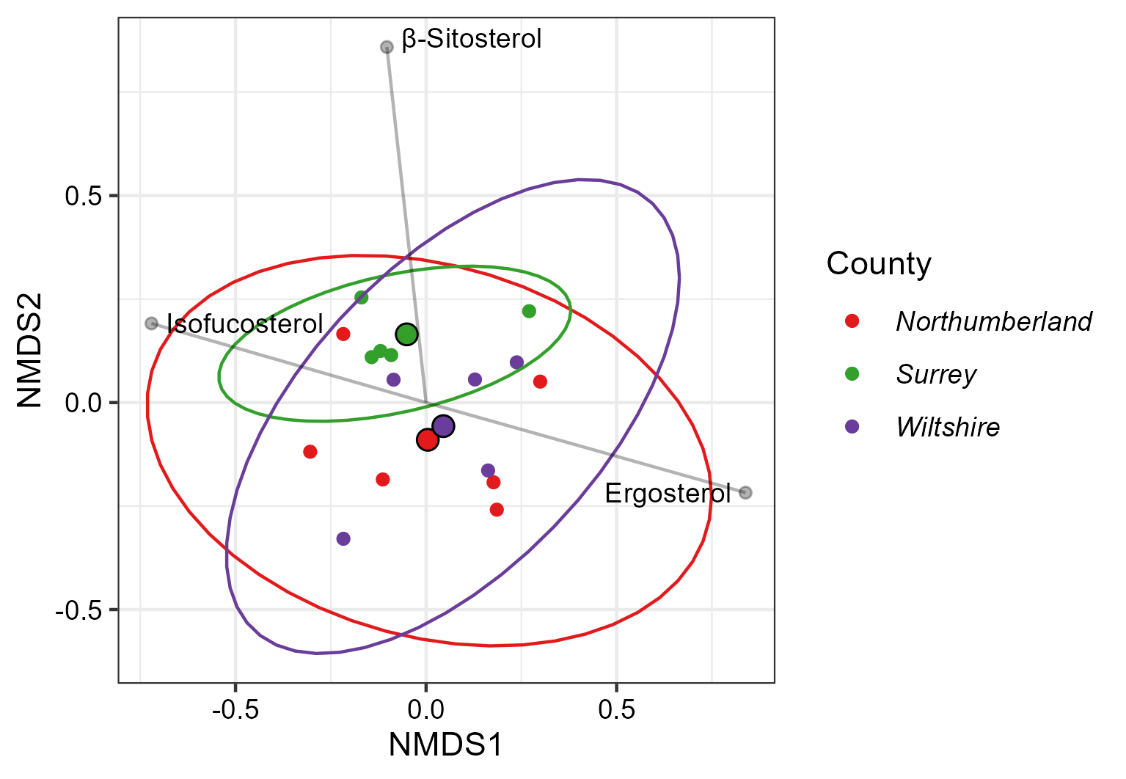

*Bombus lapidarius*

*Bombus pascuorum*

*Bombus terrestris/lucorum agg.*

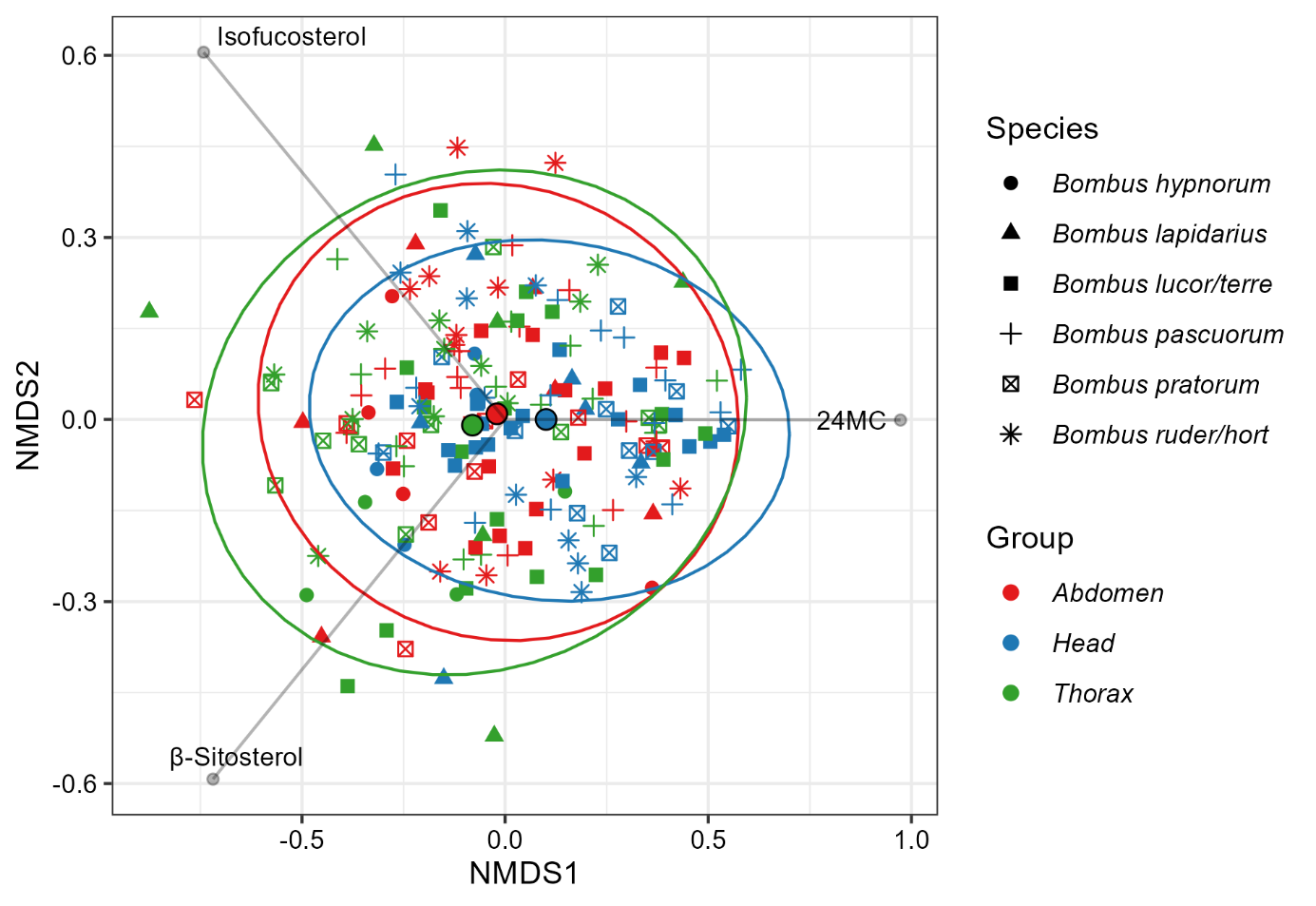

***Figure S5.*** *NMDS (stress value 0.150) comparing sterol profile of body tagmata. Results from the PERMANOVA showed that sterol profile differed significantly between tagma (F_2,181_=4.581, p=0.001) and as a result of species identity (F_5,181_=2.382, p=0.001). However, the variance explained by both was very low (Tagmata: R^2^=0.045, Species: R^2^=0.059). There was no significant difference in variance between tagmata groups (F_2,186_=2.005, p>0.100) but there was between species groups (F_5,183_=2.420, p<0.050). The results of pairwise comparisons showed that the sterol profile of the head differed significantly from the abdomen (p<0.050) and thorax (p<0.010) but there was no difference between the abdomen and thorax (p>0.100). Investigating the effect of species using pairwise comparisons showed that the sterol profile of* B. hortorum/ruderatus *samples differed significantly from* B. terrestris/lucorum *and that* B. hyponorum *differed significantly from* B. terrestris/lucorum *and* B. pascuorum *(all p<0.050).*

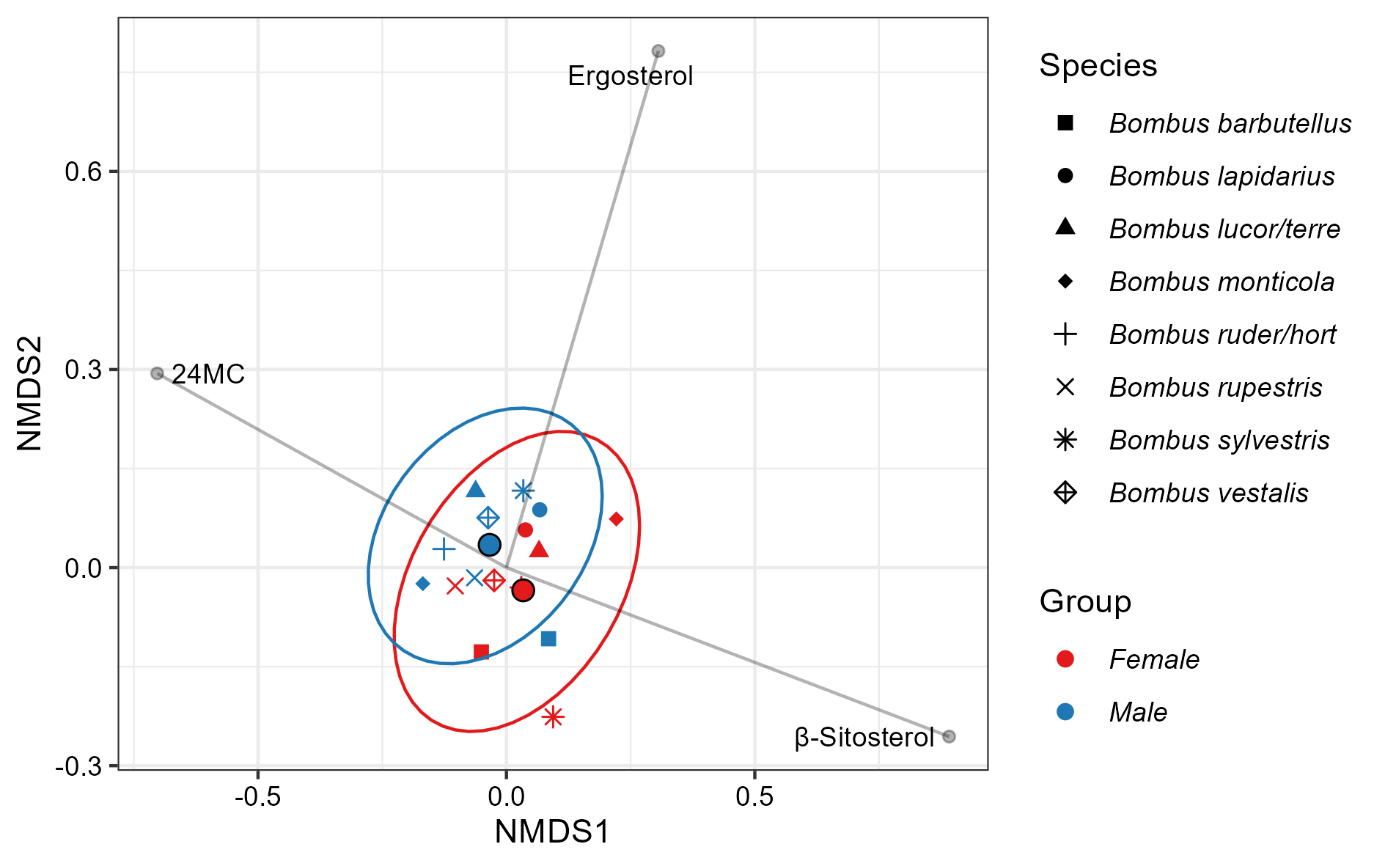

**B**

**A**

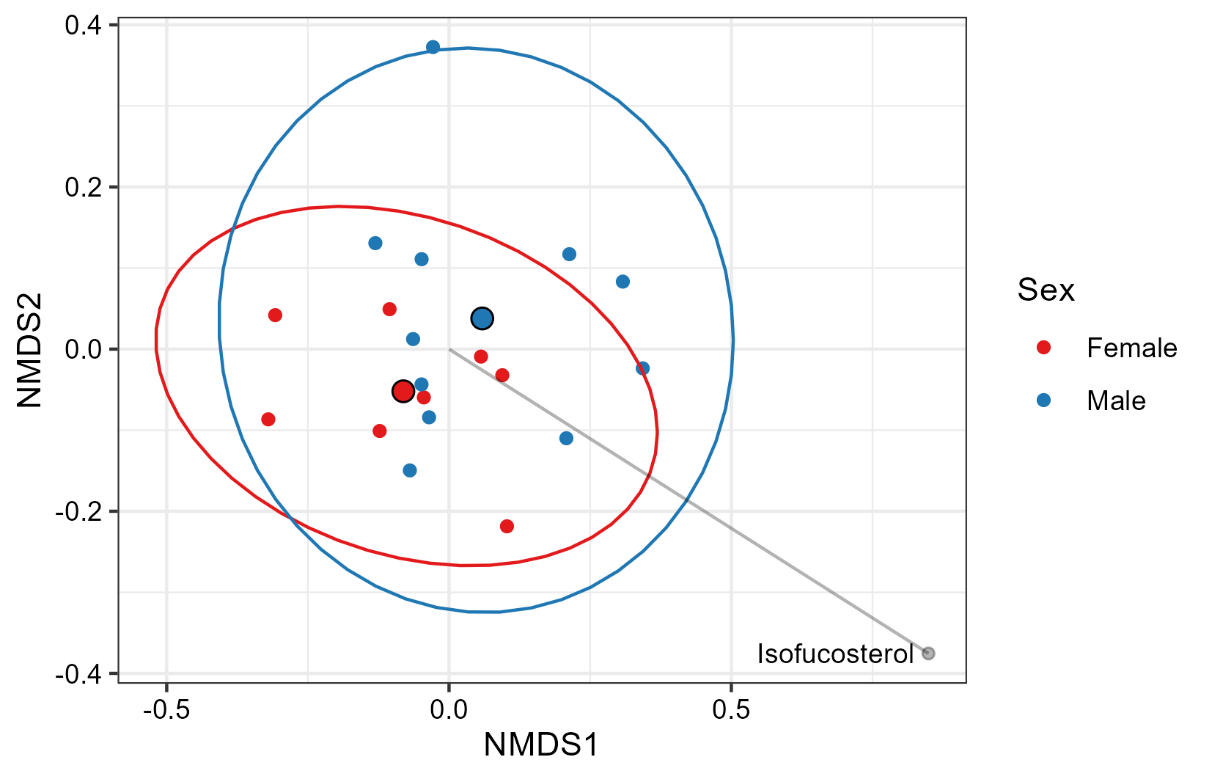

***Figure S6***

***A)*** *NMDS comparison of male and female* B. vestalis *specimens showing no significant difference between sexes. Stress value = 0.121.*

***B)*** *NMDS (stress=0.138). Comparison of mean values of male and female specimens of* B. barbutellus, B. hortorum/ruderatus, B. lapidarius, B. monticola, B. rupestris, B. sylvestris, B. terrestris/lucorum *and* B. vestalis *showing no effect of sex (PERMANOVA: F_1,7_= 1.367, p>0.100, dispersion: F_1,14_=0.025, p>0.500) or species identity (PERMANOVA: F_7,7_= 0.995, p>0.500, dispersion: F_7,8_= 1.237e^30^, p<0.001) on sterol profile.*

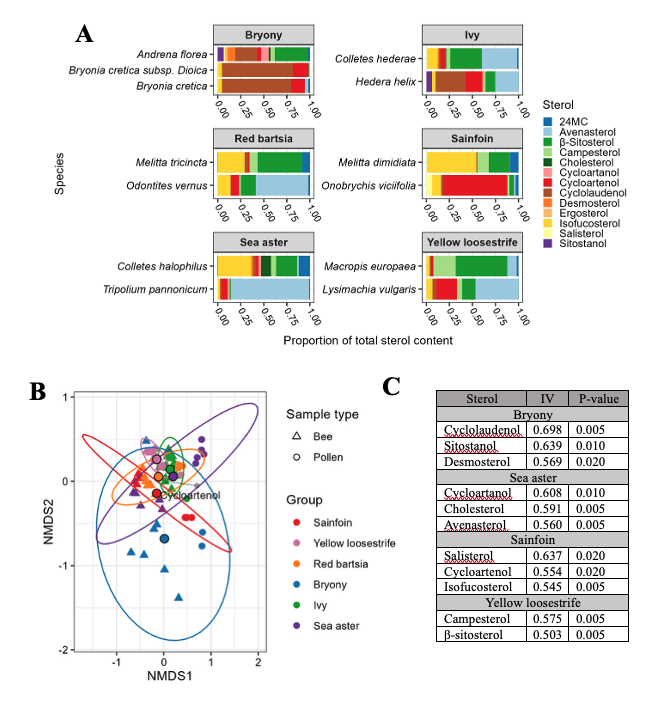

**Figure S7. *A)*** *Mean sterol profile of specialist bee species and their associated pollen. Bee species are listed first in each plot. Bars demonstrate that an abundance of a sterol in the pollen source does not guarantee its dominance in bee tissues. In general, bees contained higher proportions of β-sitosterol and isofucosterol than their pollen which contained higher avenasterol, cycloartenol and cyclolaudenol.* ***B)*** *NMDS showing monolectic and highly pollen specialist bees with their associated pollen (stress value = 0.162). Bee-pollen groups were significantly different from each other (F_5,56_=18.635, R^2^=0.401,* p*=0.001). Groups also show large differences in variation; the yellow loosestrife group and ivy group show the greatest similarity between pollen and bee sterolomes in NMDS space. In contrast, the Bryony group showed both large variation and a sterolome more distinct from the other groups. Grey lines indicate significant (*p*<0.010) and strong correlations (absolute value >0.7).* ***C)*** *Summary of Indicator Species Analysis results showing associations between groups in Figure 8B and individual sterols. Neither the ivy or red bartsia groups were associated with individual sterols, potentially due to their high overlap with other group sterolomes.*
